# Do aposematic species have larger range sizes? A case study with Neotropical poison frogs

**DOI:** 10.1101/2023.11.16.567343

**Authors:** Priscila Silveira, Fernanda Gonçalves de Sousa, Philipp Böning, Natan M. Maciel, Juliana Stropp, Stefan Lötters

## Abstract

**Aim:** Aposematic animals, i.e. those that are defended and warn potential predators through signals, are suggested to have resource-gathering advantages against non-aposematic ones. We here explore this in a biogeographic frame expecting that aposematic species are better dispersers, which translates into larger geographic range size.

**Location:** South America.

**Taxon:** Poison frogs (Amphibia; Aromobatidae and Dendrobatidae).

**Methods:** We use 43 toxic and 26 non-toxic poison frog species from the lowlands only as representatives of aposematic and non-aposematic study organisms, respectively. Realized and potential geographic ranges are calculated using minimum convex polygon and species distribution modelling methods, respectively. Accounting for species body size and phylogeny, we test if both range and aposematism are correlated using linear mixed models.

**Results:** Aposematic and non-aposematic species do neither differ in realized nor in potential geographic range size. There was no effect of body size.

**Main conclusions:** The role of aposematism yet is not as clear as suggested and determinants of poison frog range sizes are multifaceted. A more integrative approach is needed using information of behaviour, predation risk, and reproductive biology to assess the role of aposematism on observed species distributions. Such data are not yet available for most species, neither poison frogs nor other aposematic animals.

## 1 INTRODUCTION

Animals that are defended and warn potential predators through signals are called aposematic (Ruxton et al., 2004). For instance, numerous species use bold colours to caution attackers of their potent toxins, such as marine opisthobranchs, tiger moths, paper wasps, ladybeetles and poison frogs (Bezzerides et al., 2007; Cortesi & Cheney, 2010; Maan & Cummings, 2012; Nokelainen et al., 2014). Aposematism can be costly for the organism but reduces the risk of predation (Endler & Mappes, 2004; Mappes et al., 2005; Holen & Svennungsen, 2012; Stevens & Ruxton, 2012; Summers et al., 2015) and confers resource-gathering advantages (Speed et al., 2010).

In recent years, extensive work on evolutionary and functional aspects of aposematism in prey-predator relationships has become available (e.g. Mappes et al., 2005; Stevens & Ruxton 2012; Summers et al., 2015). Across vertebrates and invertebrates, aposematic animals tend to show increased movement and forage more openly, so that larger home ranges and wider niches are exploited, while camouflaged animals are associated with reduced movement and limited resource use including space (Stamp & Wilkens, 1993; Merilaita & Tullberg, 2005; Cooper et al., 2009; Mochida, 2009, Arbuckle et al., 2013). Simulation models corroborate that behavioural conspicuousness is a vital complement to aposematic signals (Speed & Ruxton, 2007; Speed et al., 2010).

Neotropical frogs of the genus *Oophaga* (family Dendrobatidae) are a well-studied example of aposematism. These amphibians exhibit remarkable within-species variation in colour and toxicity (Daly & Myers, 1967; Maan & Cummings, 2012). Signal intensity, i.e. colour conspicuousness, is generally ‘honest’ as it positively correlates with toxicity (Summers et al., 2015). As a consequence, more colourful males of the strawberry poison frog, *Oophaga pumilio*, call from more open sites to attract females and forage more openly, as the risk of predation is diminished (Pröhl & Ostrowski, 2011; Pröhl et al., 2013; Willink et al., 2013).

Yet, the link between aposematism and biogeography is poorly understood, but we expect that increased movement, larger home ranges, wider niche exploitation and more bold behaviour under a reduced predation risk might favour dispersal in aposematic animals. In general, species with a high dispersal ability have been shown to have larger geographic range sizes (e.g. Dennis et al., 2000; Böhning-Gaese et al., 2006; Luo et al., 2019). A meta-analysis by Alzate & Onstein (2022) revealed that dispersal positively affects geographic range size. However, the dispersal-range size relationship can also be affected by other proxies.

We here ask: Is aposematism a key factor determining distribution size? Neotropical poison frogs (families Aromobatidae, Dendrobatidae; Fig 1), which comprise 341 species, are an excellent group to study this. Roughly, half of all species are neither brightly coloured nor toxic (Lötters et al., 2007; Kahn et al., 2015; Grant et al., 2017). Using these amphibians, we for the first time study aposematic vs. non-aposematic species in a biogeographic context and hypothesize that aposematic species have larger geographic ranges than non-aposematic species.

**Figure 1.**
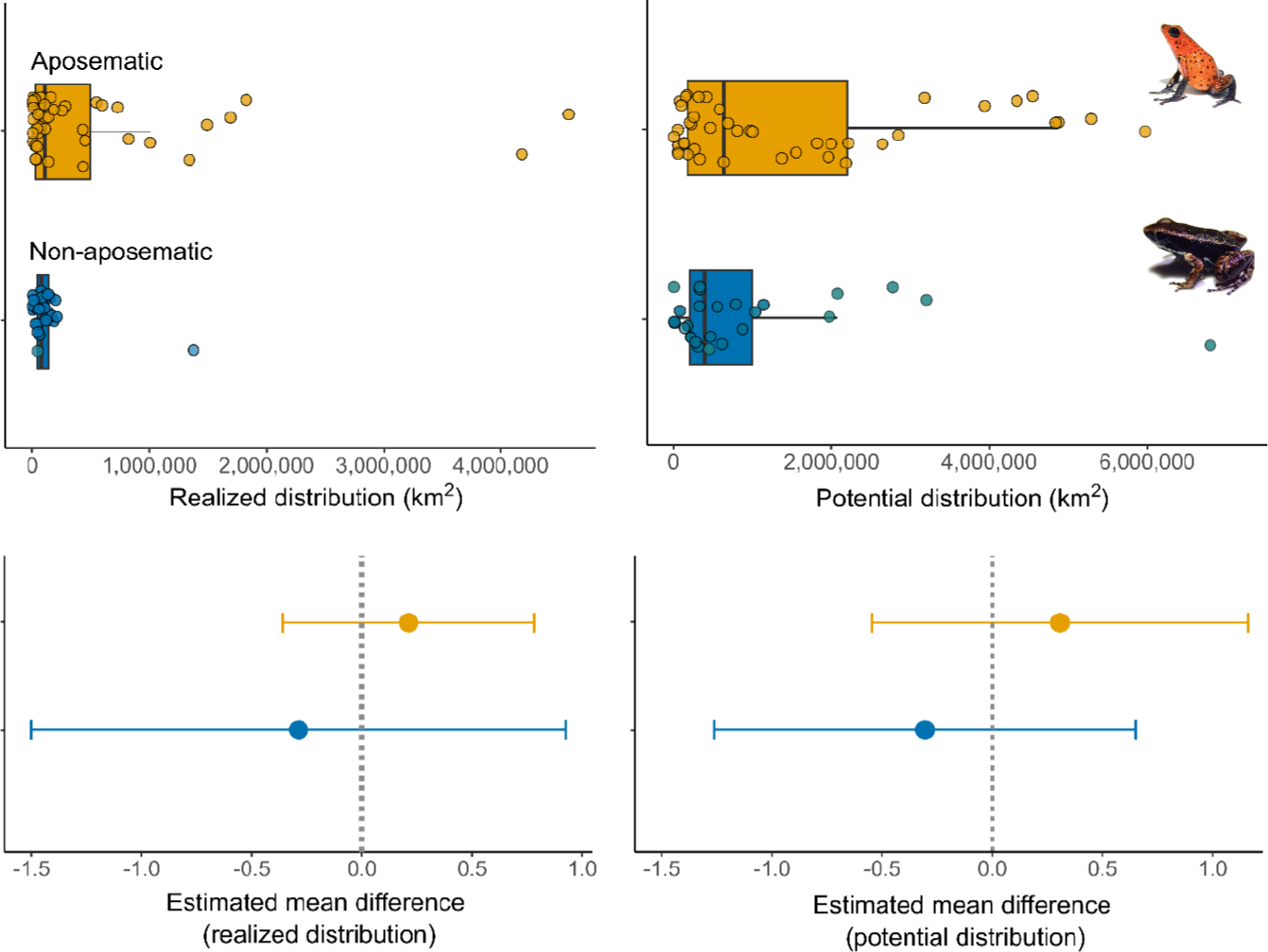
Comparison of realized and potential distribution sizes, and their mean differences in aposematic (yellow) and non-aposematic (blue) poison frog species. Aposematic and non-aposematic species are represented by *Oophaga pumilio* and *Allobates talamancae* (species photographs were courtesy of A. Plewnia).

## 2 METHODS

### 2.1 Study organisms

Poison frogs (Aromobatidae, Dendrobatidae) are distributed in northern South America and adjacent Central America. Species range from lowland rainforest, e.g. in the Amazon basin, up to tree line in the Andes (Lötters et al., 2007; Kahn et al., 2015). The high and complex mountain ranges of the Andes severely hamper species dispersal (cf. Condamine et al., 2018). For that reason, we excluded Andean taxa from our analysis. We used a set of species from lowlands only and excluded taxa that occur with at least 25% of their geographic range > 1,000 m a.s.l. This divides the species into west- and east-Andean subsets. The final data set contained 69 species belonging to the genera *Adelphobates, Allobates, Ameerega, Andinobates, Anomaloglossus, Colostethus, Dendrobates, Epipedobates, Hyloxalus, Oophaga, Phyllobates, Ranitomeya* and *Silverstoneia*. We followed the taxonomy of Grant et al. (2017) and the full list of study species is available in Table S1 in Supporting Information.

Aposematism exists along a continuum, and the degree of aposematism among (and even within) poison frog species varies. However, such continuum is hard to assess without detailed data on toxicity and colour conspicuousness, which is available for only a handful of species (Darst et al., 2006; Maan & Cummings, 2012; Willink et al., 2013). We therefore simply compare species of two groups: (i) taxa that are chemically defended and (ii) those that are not (cf. Lötters et al., 2007; Kahn et al., 2015). To allocate species to one of these two groups, we used known presence vs. absence of skin alkaloids as proxies, because with a few exceptions, as soon as bold colour signals occur in poison frogs, they are toxic (cf. Kahn et al., 2015; Summers et al., 2015). Therefore, we coded species as ‘aposematic’ (N = 43) and ‘non-aposematic’ (N = 26) when alkaloids were present and absent, respectively. With regard to the two aforementioned lowland regions, we studied 25 east-Andean aposematic, 17 east-Andean non-aposematic, 18 west-Andean aposematic and 9 west-Andean non-aposematic species (see Appendix 1 in Supporting Information; Fig. S1, Table S1).

### 2.2 Range size

We included both the realized and the potential distributions of the study organisms because the realized distribution can be affected by biotic interactions, e.g. niche competition (Peterson et al. 2011) and may suffer from sampling bias, and therefore, results could potentially be misleading. For this reason, we also examined the potential distribution derived from species distribution models (SDMs) which are robust against the issues that arise when using georeferenced occurrence records of species.

#### 2.2.1 Realized distribution

Georeferenced (latitude/longitude) occurrence records of the 69 poison frog species were obtained from GBIF (https://www.gbif.org/, accessed 7 May 2021, the literature (Silverstone, 1975; 1976; Myers & Daly, 1976; Coloma, 1995; Brown et al., 2011), from unpublished data by experts, and our own collection records. Species occurrence records were inspected and cleaned in order to eliminate duplicates and outliers. The mean number of total records per species was 58, with a range of 7–464 (Table S2 in Supporting Information). Subsequently, to represent realized distributions, we drew a minimum convex polygon (MCP) enclosing 100% of occurrence records of each species and then calculated range size in km^2^ (Table S1), using the minimum bounding geometry tool in QGIS v.3.10 (https://qgis.org/). We then excluded the polygon areas that overlapped with the sea by clipping those to the limits of continental bounds and obtained the area of species’ realized distribution.

#### 2.2.2 Potential distribution

Potential distributions were estimated based on aforementioned georeferenced species records and grid-based climatic variables taken from the CHELSA database v.1.2 (https://chelsaclimate.org/, accessed 22 March 2022) at 30-arc-seconds (∼1 km) spatial resolution (Booth et al., 2014; Karger et al., 2017) as proxies of spatial environmental information (Elith & Leathwick, 2009; Elith et al., 2010). To obtain feasible models, we accepted a minimum number of six records per species (cf. Proosdij et al., 2016). We used a thinning procedure, which retained unique occurrences that were at least two grid cells apart to reduce autocorrelation in occurrence data and sampling bias (Velazco et al., 2019). We reduced the number of original climate variables to avoid collinearity using Principal Component Analysis (PCA) to derive principal components (PC), as recommended by De Marco & Nóbrega (2018). The first six PCs accounted for ∼96% of the original climate variables’ variation (Table S3 in Supporting Information).

We used three modelling methods to predict species potential distribution: Maxent (Phillips et al., 2006; 2017), Random Forest (RDF; Breiman, 2001), and Support Vector Machine (SVM; Tax & Duin, 2004). Maxent is a presence / background method that operates with a machine-learning algorithm following the principle of maximum entropy. It is considered a reliable method and is widely used (Peterson et al. 2011). However, Maxent requires some caution, because uncritical use of settings (described by Phillips et al., 2006; Phillips & Dudík, 2008) can drastically alter and mislead the output (Elith et al., 2010; 2011; Merow et al., 2013; Yackulic et al., 2013). We therefore applied settings in relation to the available data (as recommended by Phillips & Dudík, 2008; Warton & Aarts, 2013; Radosavljevic & Anderson, 2014; Phillips et al., 2017). RDF is a machine learning method using presence / pseudo-absence information (Breiman, 2001; Peterson et al. 2011). It is a robust model and one of the most recommended algorithms for predicting species distribution (Evans et al., 2011; Liaw & Wiener, 2002). SVM employs presence / pseudo-absence information and can obtain a better data boundary than other methods, since it solves the multidimensional outlier detection problem (Tax & Duin, 2004). SVM shows comparable or better results for complex data sets when outlier information is used (Tax & Duin, 2004; Peterson et al. 2011).

A background area, which enclosed Central and South America, was used for all species as a window spanning the geographic ranges of all species studied. Then, we bootstrapped species-occurrence dataset into two subsets: 1) keeping 70% of the initial data out of the calibration for the model training validation and 2) using 30% for model testing. Additionally, we applied environmental constrains as pseudo-absence allocation method and background points based on the lowest predicted suitable region within the area used to calibrate the models (Engler et al., 2004). The use of the calibration area with the pseudo-absences performs better since the performance of the algorithms may be sensible to the way pseudo-absences are allocated (Wisz & Guisan, 2009; Andrade et al., 2020). We restricted the available climatic information to train the models with the ecoregions shapefile from World Wildlife Fund website (https://www.worldwildlife.org/biomes) and restricted the models to ecoregions with known occurrences of the study species. This approach avoids model over-fitting and unreliable predictions for the species distribution range (VanDerWal et al., 2009; Barve et al., 2011).

To avoid underestimating the species’ distribution range, we defined threshold of suitability value based on values of maximum sensitivity and specificity obtained with Jaccard Index (Leroy et al., 2018). This step avoids under-estimating species’ distributions. To ensemble the final distribution range, we used the average of suitability values, weighted by the performance of the three algorithms for all species (Thuiller et al., 2009). We used the Jaccard Index, whose values ranges from 0 to 1, as the evaluation metric to predict accuracy of each algorithm and ensemble methods (Leroy et al., 2018), considering values equal or higher than 0.7 as acceptable (Leroy et al., 2018). The binary maps of the ensemble models were used to calculate the potentially occupied geographic range of each species. We ran all SDM analyses with the “ENMTML” R package (Andrade et al., 2020; R CoreTeam, 2023). We then re-projected the raster maps of each species to SAD 69 Albers Equal Area (EPGS 102033) and estimated the area in km^2^ (Table S1) using QGIS 3.1.

### 2.3 Statistical analyses

As species are not independent samples due to shared ancestry, and closely related species tend to have similar traits (Felsenstein, 1985), it is necessary to control for the effect of phylogenetic correlation when regressing species traits (Harvey & Pagel, 1991). We used a pruned version of the consensus poison frog phylogeny from Grant et al. (2017) to construct the variance-covariance matrix and account for phylogenetic dependency of traits on our analysis. We fit linear mixed models (lmm) to test whether aposematism is correlated to variation on both the realized (MCP) and potential distribution (SDM) of poison frogs. For this, we included the two regions (east-Andes, west-Andes) as random factor since they strikingly differ in spatial extension. Subsequently, we tested whether body size of poison frogs correlated with our main response variable (aposematism) using the *gls*() function from the ‘nlme’ package (Pinheiro et al., 2020) including a phylogenetically correlated error structure in the model to perform this test. We found no significant correlation (see Appendix 1 in Supporting Information; Figure S2, Table S4) thus, we used body size (i.e. maximum recorded adult snout-vent length; Table S1 in Supporting information) as a covariate to control for its effects on realized and potential range sizes (Womack & Bell 2020, Alzate et al., 2022). Hence, our model was coded as Y∼ Aposematic + Body size + (1|Region).

Finally, we accounted for effects of shared evolutionary history by including Pagel’s lambda (λ; Pagel 1999) correlation structure in the lmm. Pagel’s λ ranges from 0, indicating no phylogenetic signal (traits evolve independently from the phylogenetic relationship of species), to 1 indicating that traits are evolved following a Brownian motion process (close relatives are more similar to each other than to distant ones; Pagel, 1999). Correlation structures were incorporated in the models using the *corPagel*() function from the “Geiger” R package (Harmon et al., 2008; Pennel et al., 2014).

Lmm was implemented through the *lme* () function available in “nlme” R package (Pinheiro et al., 2020). All statistical analyses were performed in R 4.0.2 (R CoreTeam, 2023).

## 3 RESULTS

The mean area of realized distribution of aposematic species was 515,673.5 km^2^ (± 987,676.3 km^2^), whilst that of non-aposematic species was 137,777.5 km^2^ (± 260,142.9 km^2^). The area of aposematic species’ potential distributions averaged 1,486,991 km^2^ (± 1,732,029 km^2^) and 969,060.3 km^2^ (± 1,462,787) in non-aposematic species’ with overall model accuracy (Jaccard Index) averaging 0.85 (± 0.08; Table S5). Phylogenetic signal was 1.5 times smaller in realized distribution, compared to the one estimated for potential distribution (in terms of Pagel’s lambda; Table 1). Average adult body size was 24.18 mm (±7.89; Table S1) and we found no significant correlation between body size and size of the realized or the potential distribution (Table 1).

**Table 1.** Summary statistics of the phylogenetic linear mixed models and estimated phylogenetic signal (Pagel’s lambda). Non-aposematic species were the intercept in the model, with: 0 = non-aposematic; 1 = aposematic. Model estimates of range size shown below correspond to log-transformed km^2^ values.

| Response | Pagel's $\lambda$ | Aposematic | Value | Standard error | t-value | DF | p-value |
| --- | --- | --- | --- | --- | --- | --- | --- |
| Realized range size | 0.29 | 0 (Intercept) | 5.00 | 0.40 | 12.42 | 65 | < 0.05 |
|  |  | 1 | 0.18 | 0.25 | 0.71 |  | 0.47 |
|  |  | Body size | -0.01 | 0.01 | -0.83 |  | 0.40 |
| Potential range size | 0.44 | 0 (Intercept) | 5.40 | 0.49 | 11.00 | 65 | < 0.05 |
|  |  | 1 | 0.15 | 0.20 | 0.74 |  | 0.46 |
|  |  | Body size | 0.00 | 0.00 | 0.31 |  | 0.75 |

Our study revealed that aposematic and non-aposematic species do not differ in either the realized distribution (MCP) or the potential distribution (SDM) (Table 1, Fig. 1). Additionally, potential and realized distributions showed similar mean and variance in aposematic species whilst potential distribution size of non-aposematic species showed much greater variance when compared to their realized distribution derived from MCPs (Fig 1).

## 4 DISCUSSION

### 4.1 Aposematism and geographic range size

Behavioural conspicuousness is increased in aposematic animals, and this is suggested to go in line with resource-gathering benefits (Speed & Ruxton, 2007; Speed et al., 2010). Here, we explore this rationale in a biogeographic context, expecting that aposematic species had larger geographic ranges than non-aposematic ones. Therefore, we expected that aposematic species with their bold behaviour combined with a reduced predation risk would have dispersal advantages and this would be manifested in the sizes of their realized and potential geographic ranges. Dispersal is one of the main drivers of species’ spatial distributions (Lester et al., 2007; Alzate & Onstein, 2022), and an increased dispersal ability has been demonstrated to positively influence range size in a variety of invertebrates and vertebrates such as butterflies (Dennis et al., 2000), birds (Böhning-Gaese et al., 2007; Capurucho et al., 2020), bats (Luo et al., 2019) and amphibians (Penner & Rödel 2019).

Using aposematic and non-aposematic poison frogs (Aromobatidae, Dendrobatidae), we hypothesized that aposematic species had larger geographic ranges than non-aposematic species. Our results did not support this hypothesis – there is no evidence that aposematic species have larger geographic ranges. However, does this mean that aposematism is not beneficial to dispersal and/or range expansion?

#### 4.1.1 Dispersal ability

According to Penner & Rödel (2019), who studied dispersal in West African savannah amphibians, dispersal ability correlates positively with geographic range size, suggesting that better dispersers have larger geographic range sizes. However, we find it difficult to adopt the proxies used by Penner & Rödel (2019) to poison frogs, as we do not find much interspecific trait variance (i.e. habitat and diet niche breadth) in our focal groups. We deduce that dispersal ability in aromobatids and dendrobatids might be influenced by two aspects; their complex reproductive biology as well as their food specialization (Santos et al., 2003; Lötters et al., 2007; Saporito et al., 2007; Carvajal-Castro et al., 2021). Apart from female mating choice, laying relatively few terrestrial eggs and transporting tadpoles to water bodies, which is obligate in almost all poison frogs, certain species show complex patterns of parental care behaviour including larval feeding with maternal eggs until metamorphosis is completed (e.g. in the genus *Oophaga*; Pröhl & Hödl, 1999). In several species this is even accompanied by pair bonding (e.g. in the genus *Ranitomeya*; Brown et al., 2008; 2010; Caldwell, 1997; Pettitt et al., 2020). These are characters that perhaps do not enhance dispersal but favour territoriality (e.g. Pröhl, 1997; 2005; Wells, 2007). While a more promiscuous mating behaviour (mating multiple times with several partners) may be associated to a broader home range, habitat use and, potentially, translating into higher dispersal rates (e.g. in *Allobates femoralis*, Ursprung et al., 2011), specialized reproductive biology along with territoriality (Carvajal-Castro et al., 2021), may diminish dispersal and consequently aposematic species may have smaller geographic range sizes. Surprisingly, our results show that this (aposematic poison frogs have smaller ranges) is also not the case among poison frogs in general.

Food specialization is another distinctive aspect in poison frogs, as in particular many aposematic species focus on small prey items, especially ants and mites (Santos et al., 2003; Saporito et al., 2007). However, it remains unstudied if myrmecophagy in poison frogs has such an effect because small ants and mites might be widespread in tropical lowlands (Beck, 1971). Moreover, a negative association between diet breadth and range size was reported in microhylid frogs of the genus *Coxiphalus* living in the rainforests of Queensland, Australia, where food specialists, mostly myrmecophagous like the aposematic poison frogs, had larger range sizes (Williams et al., 2006). Lastly, many poison frogs occur along streams, which often applies to non-aposematic species (e.g. *Hyloxalus, Mannophryne* or *Leucostethus*, Lötters et al., 2007; Grant et al., 2017; Table S1). Generally, a riparian lifestyle is suggested to favour long-distance dispersal in amphibians (Marin da Fonte et al., 2019) due to habitat dynamics and high connectivity of river systems. If indeed this is true, obviously a larger number of non-aposematic species would benefit from this dispersal advantage.

### 4.2 Aposematism: a phenotypic response to spatial constraints?

Although dispersal has been demonstrated to increase geographic range size in species (dispersal-range size relationship), other factors may also play a role. Such factors include organism specific traits like body size and shape, life stage (i.e. seeds, egg type, size, clutch size, larval type) and life span (Alzate & Onstein, 2022). So far, accessing the effects of life history aspects in aromobatids and dendrobatids on their dispersal abilities is premature due to the absence of robust data. However, at least we *ad hoc* find reasonable that aposematic species have dispersal disadvantages compared to non-aposematic species. Despite this, geographic ranges in these two groups do not differ in size, suggesting that hypothetically aposematism compensates dispersal disadvantages.

### 4.3 Conclusion and outlook

This study presents a first attempt to uncover the role of aposematism in a biogeographic frame, using Neotropical poison frogs. Contrary to our expectations, we found no evidence that aposematic species have wider geographic ranges than non-aposematic species. Given the data, we found no support for a strong signal favouring that aposematism is a key factor delimiting geographic range size. However, for now, our data also do not demonstrate that aposematism has no effect. We instead conclude that the connection between aposematism and distribution is far more complex and results from an interplay of aposematism – in poison frogs – with other factors such as phenotypic traits, behaviour, predation risk and reproductive biology. Yet, empirical data on each of these factors remain scattered in the scientific literature, which hampers a more integrative analysis.

Additionally, our results suggest potential limitations in occurrence data of non-aposematic species compared to aposematic species given the observed variance between the calculated potential and realized distributions in this group. This may be due to their crypsis leading to a less representative sampling, while aposematic species due to their bright coloration are easier to spot. The fact that there are more studies available on aposematic than on non-aposematic poison frogs (e.g. Lötters et al., 2007; Kahn et al., 2017) can be an explanation for the greater variance found for this group.

We call for the need of increasing knowledge on species-specific traits and distribution records in poison frogs. In combination, this will in turn not only enable better distribution modelling but also accurate predictions of how habitat loss and climate change will affect poison frogs in the Neotropics.

## Acknowledgements

We are grateful to Jason L. Brown and Evan Twomey for sharing poison frog distribution records with us. We also thank Evan for insightful comments on previous versions of the manuscript. PS was supported by Coordenação de Aperfeiçoamento de Pessoal de Nível Superior (CAPES, grant number: 88881.361853/2019-01). NMM is supported by Conselho Nacional de Desenvolvimento Científico e Tecnológico (CNPq).

## DATA AVAILABILITY STATEMENT

Data used on this study are available in XXXXXXXX

## BIOSKETCH

Priscila Silveira is broadly interested in intrinsic and extrinsic factors that shape spatial and temporal distribution of all types of organisms, especially anurans.

## SUPPORTING INFORMATION

Additional supporting information may be found online in the Supporting Information section.

## AUTHOR CONTRIBUTION

PS and SL developed the concept of this study, collected and curated the data and performed the analyses along with FGS. All authors discussed the results and contributed to the writing.

