## Supplementary Material for "Do aposematic species have larger range sizes? A case study with Neotropical poison frogs"

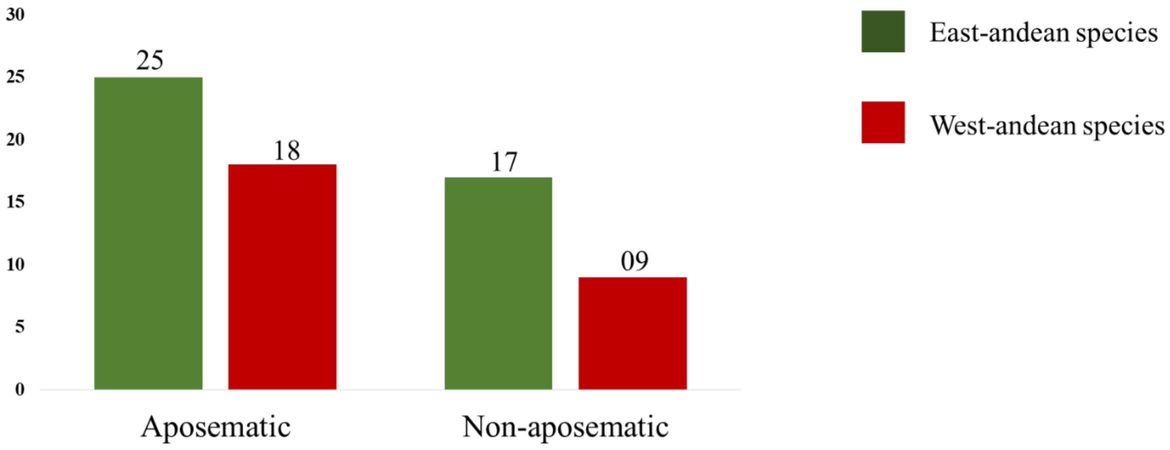


**Figure S1.** Number of species included in our study. Our dataset comprised of 25 aposematic and 17 non-aposematic species from the East side of the Andes and 18 aposematic and nine non-aposematic species occurring on the west side of the Andes.

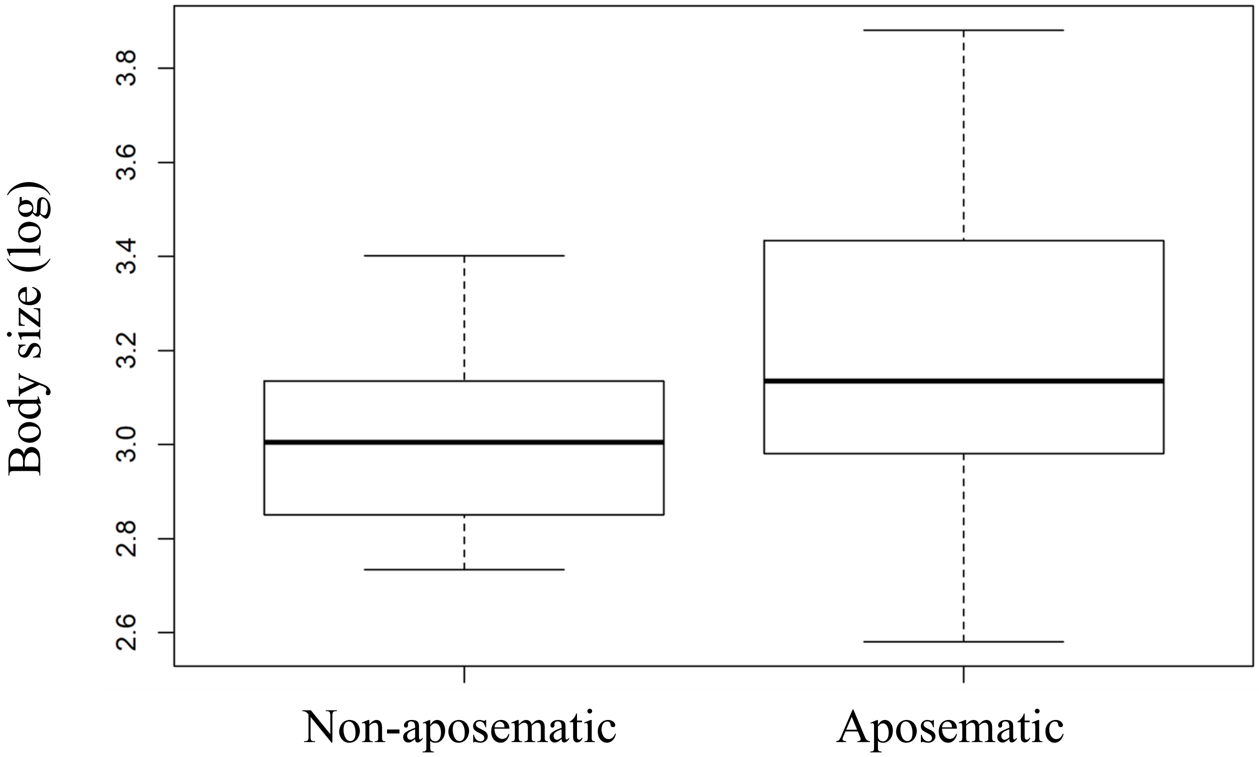


**Figure S2.** Body size in log-scale of non-aposematic and aposematic species included in our study. There is no significant difference between body sizes of aposematic and non-aposematic species ($p>0.05$).

**Table S1.** Species and their information used in the the study. Aposematic species (1); Non-aposematic species (0) and the respective absolute and logaritmized values of area size in km² of their realized and potential distribution range and maximum body size.

| Species | Region | Aposematic | area_realized_distribution_km2 | area_potential_distribution_km2 | log_potential_distribution | log_realized_distribution | maximum_body_size | log_maximum_body_size |
|---|---|---|---|---|---|---|---|---|
| Adelphobates_castaneoticu | east | 1 | 50569.3 | 170398.4321 | 5.231465594 | 4.7038869 | 22.7 | 3.122364924 |
| Adelphobates_galactonotu | east | 1 | 1005681.6 | 2181106.435 | 6.338676859 | 6.0024605 | 40.5 | 3.701301974 |
| delphobates_quinquevittat | east | 1 | 450237.5 | 4548817.368 | 6.657898501 | 5.6534417 | 16.5 | 2.803360381 |
| Allobates_cepedai | east | 0 | 5579.7 | 329092.8069 | 5.51731839 | 3.7466108 | 19.5 | 2.970414466 |
| Allobates_granti | east | 0 | 135120.8 | 615506.9976 | 5.789232995 | 5.1307222 | 16.18 | 2.783775912 |
| Allobates_kingsburyi | east | 0 | 35431.1 | 1973404.447 | 6.295216103 | 4.5493846 | 20.6 | 3.025291076 |
| Allobates_nidicola | east | 0 | 54460.2 | 1137567.488 | 6.055977171 | 4.7360792 | 19.6 | 2.975529566 |
| Allobates_niputidea | west | 0 | 128633.2 | 15260.36755 | 4.183564994 | 5.1093531 | 16.75 | 2.818398258 |
| Allobates_paleovarzensis | east | 0 | 184275.3 | 2077615.251 | 6.317565125 | 5.2654671 | 21.7 | 3.077312261 |
| Allobates_sumtuosus | east | 0 | 45636.7 | 178104.9651 | 5.250676027 | 4.6593142 | 15.4 | 2.734367509 |
| Allobates_talamancae | west | 0 | 163735.8 | 335947.0521 | 5.526270835 | 5.2141436 | 21.9 | 3.086486637 |
| Allobates_tapajos | east | 0 | 103278.7 | 447687.6941 | 5.650975157 | 5.0140108 | 15.9 | 2.766319109 |
| Allobates_trilineatus | east | 0 | 1376925.3 | 2777285.394 | 6.44362051 | 6.1389104 | 17.09 | 2.838493497 |
| Allobates_zaparo | east | 0 | 73763.4 | 6799054.956 | 6.832448552 | 4.8678409 | 27.9 | 3.328626689 |
| Ameerega_bassleri | east | 1 | 3435.8 | 638070.412 | 5.804868606 | 3.5360279 | 42 | 3.737669618 |
| Ameerega_bilinguis | east | 1 | 82531.8 | 977674.8565 | 5.990194446 | 4.9166213 | 20.95 | 3.042138646 |
| Ameerega_braccata | east | 1 | 432731.4 | 463032.3836 | 5.665611366 | 5.6362184 | 21.8 | 3.08190997 |
| Ameerega_flavopicta | east | 1 | 1826551 | 4348914.472 | 6.638380867 | 6.2616318 | 30.5 | 3.417726684 |
| Ameerega_hahneli | east | 1 | 4580669.7 | 4876405.892 | 6.688099847 | 6.660929 | 23 | 3.135494216 |
| Ameerega_macero | east | 1 | 133400.6 | 3178125.173 | 6.502170998 | 5.1251578 | 29.5 | 3.384390263 |
| Ameerega_parvula | east | 1 | 110489.7 | 1552719.637 | 6.191093046 | 5.0433218 | 22.5 | 3.113515309 |
| Ameerega_petersi | east | 1 | 281361.6 | 5973068.384 | 6.776197487 | 5.4492648 | 31 | 3.433987204 |
| Ameerega_picta | east | 1 | 1346271.2 | 1995468.38 | 6.30004485 | 6.1291326 | 24 | 3.17805383 |

| | | | | | | | | |
|---|---|---|---|---|---|---|---|---|
| Ameerega_trivittata | east | 1 | 4175373.1 | 4841583.07 | 6.684987388 | 6.6206953 | 48.5 | 3.881563798 |
| Andinobates_fulguritus | west | 1 | 68477.1 | 268000.364 | 5.428135384 | 4.8355454 | 15 | 2.708050201 |
| Andinobates_minutus | west | 1 | 57217.7 | 79006.51813 | 4.897662923 | 4.7575304 | 13.2 | 2.58021683 |
| ıomaloglossus_baeobatracl | east | 0 | 151430.8 | 1035669.811 | 6.015221317 | 5.1802142 | 18 | 2.890371758 |
| Anomaloglossus_beebei | east | 0 | 119284.9 | 554666.0794 | 5.744031607 | 5.0765855 | 17 | 2.833213344 |
| .nomaloglossus_degranvill | east | 0 | 209901.9 | 786472.4544 | 5.895683516 | 5.3220164 | 17.3 | 2.850706502 |
| Anomaloglossus_kaiei | east | 0 | 3323.9 | 10070.52574 | 4.003052144 | 3.5216479 | 19.8 | 2.985681938 |
| Anomaloglossus_praderioi | east | 0 | 1588.8 | 4670.449173 | 3.66935865 | 3.2010692 | 22.7 | 3.122364924 |
| Anomaloglossus_stepheni | east | 0 | 199320.6 | 3201034.639 | 6.505290374 | 5.2995522 | 18 | 2.890371758 |
| Colostethus_fraterdanieli | west | 0 | 61913.5 | 464098.7113 | 5.666610363 | 4.7917854 | 22.38 | 3.108167703 |
| Colostethus_inguinalis | west | 0 | 82151.4 | 78998.31715 | 4.89761784 | 4.914615 | 30 | 3.401197382 |
| Colostethus_panamansis | west | 1 | 29267.3 | 153500.0417 | 5.186108498 | 4.4663827 | 23.2 | 3.144152279 |
| Dendrobates_auratus | west | 1 | 134264.1 | 689416.2997 | 5.838481547 | 5.1279599 | 33.5 | 3.511545439 |
| Dendrobates_leucomelas | east | 1 | 547050.1 | 5288846.329 | 6.723360948 | 5.7380271 | 30.99 | 3.433664572 |
| Dendrobates_tinctorius | east | 1 | 593296.4 | 1816911.462 | 6.259333765 | 5.7732717 | 45 | 3.80666249 |
| Dendrobates_truncatus | west | 1 | 255349.8 | 331554.6118 | 5.520555073 | 5.4071355 | 25.6 | 3.242592351 |
| Epipedobates_anthonyi | west | 1 | 19889 | 52818.3562 | 4.722784881 | 4.2986129 | 26.5 | 3.277144733 |
| pipedobates_darwinwallac | west | 1 | 8026.3 | 184084.7444 | 5.265017799 | 3.9045154 | 19.5 | 2.970414466 |
| Epipedobates_machalilla | west | 1 | 41194.2 | 414250.747 | 5.6172633 | 4.6148361 | 17.6 | 2.867898902 |
| Epipedobates_tricolor | west | 1 | 35445 | 226801.144 | 5.355645241 | 4.549555 | 23 | 3.135494216 |
| Hyloxalus_awa | west | 0 | 32161.4 | 140075.0515 | 5.146360791 | 4.5073349 | 25.9 | 3.254242969 |
| Hyloxalus_infraguttatus | west | 0 | 114225.6 | 213003.8298 | 5.328387412 | 5.0577634 | 23.4 | 3.152736022 |
| Hyloxalus_nexipus | east | 0 | 46809.8 | 873762.5741 | 5.941393439 | 4.6703368 | 23 | 3.135494216 |
| Hyloxalus_sauli | east | 0 | 47669.3 | 217563.2145 | 5.337585467 | 4.6782388 | 28.7 | 3.356897123 |
| Hyloxalus_toachi | west | 0 | 13732.2 | 314383.4179 | 5.497459631 | 4.1377401 | 28.2 | 3.339321978 |
| Oophaga_granulifera | west | 1 | 21250.1 | 58379.4349 | 4.766259887 | 4.327361 | 22 | 3.091042453 |
| Oophaga_histrionica | west | 1 | 154537.2 | 199970.8461 | 5.300966684 | 5.189033 | 38 | 3.63758616 |
| Oophaga_lehmanni | west | 1 | 342 | 58904.29707 | 4.770146978 | 2.5340261 | 35.3 | 3.563882964 |
| Oophaga_pumilio | west | 1 | 106705.3 | 318983.3428 | 5.503768005 | 5.028186 | 25 | 3.218875825 |
| Oophaga_sylvatica | west | 1 | 29398 | 138930.1959 | 5.142796648 | 4.4683178 | 38 | 3.63758616 |
| Phyllobates_aurotaenia | west | 1 | 24548.4 | 129490.8779 | 5.112239175 | 4.3900232 | 34 | 3.526360525 |

| | | | | | | | | |
|---|---|---|---|---|---|---|---|---|
| Phyllobates_lugubris | west | 1 | 40339.7 | 58194.91304 | 4.764885024 | 4.6057327 | 24 | 3.17805383 |
| Phyllobates_terribilis | west | 1 | 1517.2 | 5463.487137 | 3.737469925 | 3.1810428 | 41 | 3.713572067 |
| Phyllobates_vitattus | west | 1 | 3334.9 | 96741.11875 | 4.985611105 | 3.5230828 | 31 | 3.433987204 |
| Ranitomeya_amazonica | east | 1 | 1491787.2 | 2645195.319 | 6.422457745 | 6.1737069 | 17.5 | 2.862200881 |
| Ranitomeya_fantastica | east | 1 | 10058.1 | 2846016.295 | 6.454237382 | 4.0025159 | 23 | 3.135494216 |
| Ranitomeya_imitator | east | 1 | 8218.8 | 1372318.222 | 6.13745483 | 3.9148084 | 22 | 3.091042453 |
| Ranitomeya_reticulata | east | 1 | 103002.5 | 804118.5971 | 5.905320106 | 5.0128478 | 17 | 2.833213344 |
| Ranitomeya_sirensis | east | 1 | 430866 | 3938533.669 | 6.595334562 | 5.6343422 | 19.9 | 2.990719732 |
| Ranitomeya_toraro | east | 1 | 729981.5 | 1003479.484 | 6.001508498 | 5.8633119 | 16 | 2.772588722 |
| Ranitomeya_uakarii | east | 1 | 1694466.7 | 2214111.443 | 6.345199476 | 6.229033 | 16.2 | 2.785011242 |
| Ranitomeya_vanzolinii | east | 1 | 50973.5 | 257922.8664 | 5.411489847 | 4.7073445 | 17.85 | 2.882003508 |
| Ranitomeya_variabilis | east | 1 | 820383.7 | 1961063.321 | 6.292491617 | 5.914017 | 21 | 3.044522438 |
| Ranitomeya_ventrimaculata | east | 1 | 183468.8 | 582235.1546 | 5.765098424 | 5.2635622 | 15.55 | 2.744060639 |
| Silverstoneia_flotator | west | 0 | 58601.5 | 281536.0672 | 5.44953404 | 4.7679087 | 18 | 2.890371758 |
| Silverstoneia_nubicola | west | 0 | 133258.9 | 333034.0671 | 5.522488661 | 5.1246962 | 23 | 3.135494216 |

**Table S2.** Number of total and thinned occurence records per species and their respective occurence region. Thinned occurences correspond to the number of unique occurences that are at least two cells appart and were kept to model species potential geographic distribution. East= east of the Andes; West= West of the Andes.

| | | Number of occurences per species | |
|---|---|---|---|
| **Species** | **Region** | **Total** | **Thinned** |
| Adelphobates_castaneoticus | East | 7 | 7 |
| Adelphobates_galactonotus | East | 14 | 14 |
| Adelphobates_quinquevittatus | East | 8 | 8 |
| Allobates_cepedai | East | 15 | 11 |
| Allobates_granti | East | 52 | 49 |
| Allobates_kingsburyi | East | 9 | 9 |
| Allobates_nidicola | East | 12 | 9 |
| Allobates_niputidea | West | 30 | 28 |
| Allobates_paleovarzensis | East | 17 | 12 |
| Allobates_sumtuosus | East | 9 | 6 |
| Allobates_talamancae | West | 113 | 97 |
| Allobates_tapajos | East | 11 | 10 |
| Allobates_trilineatus | East | 34 | 33 |
| Allobates_zaparo | East | 36 | 30 |
| Ameerega_bassleri | East | 15 | 12 |
| Ameerega_bilinguis | East | 56 | 40 |
| Ameerega_braccata | East | 9 | 9 |
| Ameerega_flavopicta | East | 19 | 18 |
| Ameerega_hahneli | East | 106 | 100 |
| Ameerega_macero | East | 13 | 13 |
| Ameerega_parvula | East | 64 | 54 |
| Ameerega_petersi | East | 13 | 12 |
| Ameerega_picta | East | 38 | 37 |
| Ameerega_trivittata | East | 206 | 165 |
| Andinobates_fulguritus | West | 22 | 19 |
| Andinobates_minutus | West | 42 | 38 |
| Anomaloglossus_baeobatrachus | East | 103 | 88 |
| Anomaloglossus_beebei | East | 19 | 16 |
| Anomaloglossus_degranvillei | East | 65 | 58 |
| Anomaloglossus_kaiei | East | 8 | 7 |
| Anomaloglossus_praderioi | East | 6 | 6 |
| Anomaloglossus_stepheni | East | 16 | 13 |
| Colostethus_fraterdanieli | West | 128 | 111 |
| Colostethus_inguinalis | West | 38 | 31 |
| Colostethus_panamansis | West | 27 | 23 |
| Dendrobates_auratus | West | 464 | 335 |
| Dendrobates_leucomelas | East | 200 | 189 |
| Dendrobates_tinctorius | East | 221 | 183 |
| Dendrobates_truncatus | West | 27 | 25 |
| Epipedobates_anthonyi | West | 64 | 56 |

| | | | |
|---|---|---|---|
| Epipedobates_darwinwallacei | West | 17 | 12 |
| Epipedobates_machalilla | West | 25 | 23 |
| Epipedobates_tricolor | West | 38 | 33 |
| Hyloxalus_awa | West | 29 | 26 |
| Hyloxalus_infraguttatus | West | 58 | 49 |
| Hyloxalus_nexipus | East | 30 | 24 |
| Hyloxalus_sauli | East | 18 | 17 |
| Hyloxalus_toachi | West | 7 | 7 |
| Oophaga_granulifera | West | 54 | 49 |
| Oophaga_histrionica | West | 91 | 86 |
| Oophaga_lehmanni | West | 18 | 11 |
| Oophaga_pumilio | West | 399 | 275 |
| Oophaga_sylvatica | West | 67 | 62 |
| Phyllobates_aurotaenia | West | 21 | 21 |
| Phyllobates_lugubris | West | 35 | 31 |
| Phyllobates_terribilis | West | 6 | 6 |
| Phyllobates_vitattus | West | 52 | 46 |
| Ranitomeya_amazonica | East | 101 | 88 |
| Ranitomeya_fantastica | East | 21 | 17 |
| Ranitomeya_imitator | East | 62 | 40 |
| Ranitomeya_reticulata | East | 39 | 34 |
| Ranitomeya_sirensis | East | 50 | 43 |
| Ranitomeya_toraro | East | 30 | 29 |
| Ranitomeya_uakarii | East | 54 | 44 |
| Ranitomeya_vanzolinii | East | 11 | 10 |
| Ranitomeya_variabilis | East | 152 | 122 |
| Ranitomeya_ventrimaculata | East | 27 | 25 |
| Silverstoneia_flotator | West | 71 | 56 |
| Silverstoneia_nubicola | West | 63 | 57 |

**Table S3**. Principal component analysis (PCA) of climatic variables used to model species potential dsitribution. The column 'Variance' corresponds to the amount of variance explained by each principal component (PC). The columns 'Bio' correspond to each climatic variable downloaded from the Chelsa database and their respective correlation to each PC. Comp= axis component.

| **Axis** | **Variance** | **Bio_01** | **Bio_02** | **Bio_03** | **Bio_04** | **Bio_05** | **Bio_06** | **Bio_07** | **Bio_08** | **Bio_09** | **Bio_10** | **Bio_11** | **Bio_12** | **Bio_13** | **Bio_14** | **Bio_15** | **Bio_16** | **Bio_17** | **Bio_18** | **Bio_19** |
|---|---|---|---|---|---|---|---|---|---|---|---|---|---|---|---|---|---|---|---|---|
| Comp.1 | 0.50 | 0.26 | -0.28 | 0.09 | -0.21 | 0.20 | 0.30 | -0.26 | 0.24 | 0.28 | 0.24 | 0.28 | 0.25 | 0.24 | 0.18 | -0.21 | 0.24 | 0.18 | 0.09 | 0.21 |
| Comp.2 | 0.26 | 0.26 | 0.10 | -0.29 | 0.17 | 0.34 | 0.17 | 0.17 | 0.27 | 0.22 | 0.30 | 0.21 | -0.22 | -0.13 | -0.30 | 0.19 | -0.12 | -0.30 | -0.28 | -0.10 |
| Comp.3 | 0.08 | 0.08 | 0.18 | -0.29 | 0.41 | 0.12 | -0.07 | 0.28 | 0.19 | -0.04 | 0.12 | -0.02 | 0.11 | -0.04 | 0.30 | -0.24 | -0.03 | 0.29 | 0.48 | -0.28 |
| Comp.4 | 0.07 | 0.07 | -0.16 | 0.30 | -0.20 | -0.04 | 0.10 | -0.23 | 0.09 | 0.06 | 0.03 | 0.09 | -0.27 | -0.49 | 0.11 | -0.23 | -0.49 | 0.10 | 0.06 | -0.36 |
| Comp.5 | 0.03 | 0.00 | -0.19 | -0.19 | -0.28 | -0.10 | -0.01 | -0.12 | 0.02 | -0.07 | -0.05 | 0.03 | 0.05 | 0.21 | -0.36 | -0.05 | 0.22 | -0.34 | 0.39 | -0.57 |
| Comp.6 | 0.02 | 0.09 | 0.38 | 0.62 | -0.09 | 0.12 | 0.01 | 0.14 | 0.10 | 0.05 | 0.07 | 0.09 | 0.01 | 0.09 | 0.01 | 0.51 | 0.08 | 0.01 | 0.31 | -0.13 |
| Comp.7 | 0.01 | 0.01 | 0.45 | 0.22 | -0.27 | 0.16 | -0.06 | 0.31 | -0.08 | 0.01 | -0.03 | 0.05 | 0.08 | 0.08 | -0.12 | -0.66 | 0.09 | -0.11 | -0.25 | -0.09 |
| Comp.8 | 0.01 | 0.02 | -0.09 | 0.32 | 0.42 | -0.05 | 0.01 | -0.09 | 0.11 | -0.04 | 0.05 | -0.07 | -0.11 | -0.05 | -0.37 | -0.32 | -0.05 | -0.33 | 0.39 | 0.42 |
| Comp.9 | 0.01 | 0.01 | -0.15 | 0.33 | 0.52 | -0.12 | 0.01 | -0.17 | 0.14 | -0.02 | 0.05 | -0.11 | -0.07 | 0.29 | 0.05 | -0.07 | 0.22 | 0.06 | -0.43 | -0.44 |
| Comp.10 | 0.00 | 0.02 | -0.04 | 0.08 | 0.04 | -0.09 | -0.08 | 0.02 | 0.30 | -0.15 | -0.02 | -0.03 | 0.82 | -0.31 | -0.19 | 0.08 | -0.15 | -0.05 | -0.16 | -0.05 |
| Comp.11 | 0.00 | 0.19 | 0.00 | -0.04 | -0.25 | -0.24 | -0.23 | 0.09 | 0.66 | -0.45 | -0.05 | 0.03 | -0.30 | 0.11 | 0.07 | -0.01 | 0.06 | 0.05 | -0.08 | 0.16 |
| Comp.12 | 0.00 | 0.01 | 0.29 | -0.11 | -0.01 | -0.38 | -0.19 | -0.17 | 0.29 | 0.72 | -0.25 | -0.14 | 0.00 | 0.03 | -0.04 | -0.02 | -0.03 | 0.01 | 0.02 | 0.01 |
| Comp.13 | 0.00 | 0.04 | -0.32 | 0.10 | -0.03 | -0.06 | -0.24 | 0.35 | -0.03 | 0.21 | 0.03 | 0.02 | -0.09 | -0.53 | 0.10 | 0.00 | 0.59 | -0.07 | -0.01 | -0.01 |
| Comp.14 | 0.00 | 0.19 | 0.49 | -0.14 | 0.06 | -0.14 | 0.19 | -0.52 | -0.10 | -0.25 | 0.10 | 0.04 | -0.03 | -0.36 | -0.03 | -0.02 | 0.40 | 0.05 | -0.01 | 0.00 |
| Comp.15 | 0.00 | 0.30 | -0.06 | 0.02 | 0.00 | -0.26 | -0.25 | 0.10 | -0.26 | 0.03 | 0.28 | 0.15 | -0.03 | 0.08 | -0.52 | 0.00 | -0.07 | 0.56 | 0.00 | 0.00 |
| Comp.16 | 0.00 | 0.30 | 0.04 | 0.00 | 0.04 | -0.37 | -0.27 | -0.01 | -0.25 | 0.00 | 0.36 | 0.23 | 0.12 | 0.12 | 0.41 | -0.01 | -0.19 | -0.46 | -0.02 | 0.01 |
| Comp.17 | 0.00 | 0.74 | -0.08 | 0.02 | -0.05 | 0.15 | 0.08 | 0.05 | -0.14 | -0.01 | -0.29 | -0.55 | 0.03 | 0.02 | 0.05 | 0.02 | -0.05 | -0.05 | 0.00 | -0.01 |
| Comp.18 | 0.00 | 0.22 | -0.03 | 0.00 | 0.22 | 0.01 | -0.01 | 0.03 | -0.09 | -0.05 | -0.68 | 0.66 | 0.01 | 0.00 | 0.00 | 0.01 | -0.01 | 0.00 | 0.00 | -0.01 |
| Comp.19 | 0.00 | 0.00 | 0.00 | 0.00 | 0.00 | 0.55 | -0.73 | -0.41 | 0.00 | 0.00 | 0.00 | 0.00 | 0.00 | 0.00 | 0.00 | 0.00 | 0.00 | 0.00 | 0.00 | 0.00 |

**Table S4**. Model summary statistics of the phylogenetic generalized least squares model between body size (Y) and aposematism. Non-aposematic species (0) correspond to the intercept of the model. 1 = Aposematic species; λ=lambda

| | Value | Std. Error | t-value | p-value | Pagel's λ |
|---|---|---|---|---|---|
| Intercept (0) | 3.08 | 0.11 | 27.64 | 0.00 | 1.30 |
| 1 | 0.11 | 0.13 | 0.89 | 0.38 | |

**Table S5.** Overall model accuracy metrics of potential distribution models for each species in the study. MXS (Maxent); RDF (Random Forest); SVM (Support vector machine).

| Species | Algorithm | AUC | Kappa | TSS | Jaccard | Sorensen | Fpb | OR | Boyce | AUC_SD | Kappa_SD | TSS_SD | Jaccard_SD | Sorensen_SD | Fpb_SD | OR_SD | Boyce_SD |
|---|---|---|---|---|---|---|---|---|---|---|---|---|---|---|---|---|---|
| Adelphobatescastaneoticus | MXS | 0.95 | 0.95 | 0.95 | 0.97 | 0.98 | 1.93 | 0.00 | 1.00 | 0.16 | 0.16 | 0.16 | 0.11 | 0.06 | 0.21 | 0.00 | 0.00 |
| Adelphobatescastaneoticus | RDF | 0.88 | 0.85 | 0.85 | 0.90 | 0.94 | 1.80 | 0.00 | 1.00 | 0.21 | 0.24 | 0.24 | 0.16 | 0.10 | 0.32 | 0.00 | 0.00 |
| Adelphobatescastaneoticus | SVM | 0.85 | 0.80 | 0.80 | 0.88 | 0.93 | 1.77 | 0.00 | 0.94 | 0.32 | 0.35 | 0.35 | 0.19 | 0.12 | 0.39 | 0.00 | 0.17 |
| Adelphobatescastaneoticus | Ensemble | 0.90 | 0.85 | 0.85 | 0.90 | 0.94 | 1.80 | 0.00 | 0.95 | 0.17 | 0.24 | 0.24 | 0.16 | 0.10 | 0.32 | 0.00 | 0.16 |
| Adelphobatesgalactonotus | MXS | 0.79 | 0.68 | 0.68 | 0.76 | 0.86 | 1.52 | 0.03 | 0.62 | 0.11 | 0.17 | 0.17 | 0.10 | 0.06 | 0.21 | 0.08 | 0.33 |
| Adelphobatesgalactonotus | RDF | 0.80 | 0.68 | 0.68 | 0.76 | 0.86 | 1.52 | 0.00 | 0.63 | 0.13 | 0.12 | 0.12 | 0.06 | 0.04 | 0.13 | 0.00 | 0.34 |
| Adelphobatesgalactonotus | SVM | 0.76 | 0.58 | 0.58 | 0.70 | 0.82 | 1.40 | 0.03 | 0.30 | 0.10 | 0.12 | 0.12 | 0.06 | 0.04 | 0.12 | 0.08 | 0.10 |
| Adelphobatesgalactonotus | Ensemble | 0.82 | 0.70 | 0.70 | 0.77 | 0.87 | 1.54 | 0.05 | 0.53 | 0.10 | 0.16 | 0.16 | 0.10 | 0.06 | 0.20 | 0.11 | 0.28 |
| delphobatesquinquevittatu | MXS | 0.65 | 0.30 | 0.30 | 0.65 | 0.77 | 1.30 | 0.00 | 1.00 | 0.24 | 0.48 | 0.48 | 0.24 | 0.16 | 0.48 | 0.00 | NA |
| delphobatesquinquevittatu | RDF | 0.95 | 0.90 | 0.90 | 0.93 | 0.96 | 1.87 | 0.00 | 1.00 | 0.11 | 0.21 | 0.21 | 0.14 | 0.08 | 0.28 | 0.00 | 0.00 |
| delphobatesquinquevittatu | SVM | 0.93 | 0.85 | 0.85 | 0.90 | 0.94 | 1.80 | 0.00 | 1.00 | 0.12 | 0.24 | 0.24 | 0.16 | 0.10 | 0.32 | 0.00 | 0.00 |
| delphobatesquinquevittatu | Ensemble | 0.93 | 0.85 | 0.85 | 0.90 | 0.94 | 1.80 | 0.00 | 0.90 | 0.12 | 0.24 | 0.24 | 0.16 | 0.10 | 0.32 | 0.00 | 0.32 |
| Allobatescepedai | MXS | 1.00 | 1.00 | 1.00 | 1.00 | 1.00 | 2.00 | 0.00 | 1.00 | 0.00 | 0.00 | 0.00 | 0.00 | 0.00 | 0.00 | 0.00 | 0.00 |
| Allobatescepedai | RDF | 1.00 | 1.00 | 1.00 | 1.00 | 1.00 | 2.00 | 0.00 | 1.00 | 0.00 | 0.00 | 0.00 | 0.00 | 0.00 | 0.00 | 0.00 | 0.00 |
| Allobatescepedai | SVM | 0.97 | 0.97 | 0.97 | 0.97 | 0.98 | 1.93 | 0.03 | 0.90 | 0.11 | 0.11 | 0.11 | 0.11 | 0.06 | 0.21 | 0.11 | 0.32 |
| Allobatescepedai | Ensemble | 1.00 | 1.00 | 1.00 | 1.00 | 1.00 | 2.00 | 0.00 | 1.00 | 0.00 | 0.00 | 0.00 | 0.00 | 0.00 | 0.00 | 0.00 | 0.00 |
| Allobatesgranti | MXS | 0.89 | 0.72 | 0.72 | 0.77 | 0.87 | 1.53 | 0.09 | 0.46 | 0.03 | 0.08 | 0.08 | 0.06 | 0.04 | 0.11 | 0.08 | 0.23 |
| Allobatesgranti | RDF | 0.96 | 0.84 | 0.84 | 0.86 | 0.92 | 1.71 | 0.07 | 0.57 | 0.04 | 0.09 | 0.09 | 0.07 | 0.04 | 0.15 | 0.05 | 0.14 |
| Allobatesgranti | SVM | 0.93 | 0.82 | 0.82 | 0.84 | 0.91 | 1.68 | 0.05 | 0.51 | 0.03 | 0.04 | 0.04 | 0.04 | 0.02 | 0.08 | 0.05 | 0.17 |
| Allobatesgranti | Ensemble | 0.92 | 0.79 | 0.79 | 0.83 | 0.90 | 1.65 | 0.03 | 0.56 | 0.03 | 0.08 | 0.08 | 0.06 | 0.04 | 0.12 | 0.05 | 0.15 |
| Allobateskingsburyi | MXS | 0.49 | 0.23 | 0.23 | 0.58 | 0.73 | 1.15 | 0.03 | 0.53 | 0.23 | 0.32 | 0.32 | 0.11 | 0.08 | 0.21 | 0.11 | 0.37 |
| Allobateskingsburyi | RDF | 0.53 | 0.33 | 0.33 | 0.59 | 0.74 | 1.18 | 0.10 | 0.65 | 0.20 | 0.27 | 0.27 | 0.07 | 0.06 | 0.14 | 0.16 | 0.30 |
| Allobateskingsburyi | SVM | 0.59 | 0.40 | 0.40 | 0.62 | 0.77 | 1.24 | 0.03 | 0.61 | 0.14 | 0.14 | 0.14 | 0.05 | 0.04 | 0.10 | 0.11 | 0.28 |
| Allobateskingsburyi | Ensemble | 0.60 | 0.43 | 0.43 | 0.64 | 0.78 | 1.27 | 0.03 | 0.51 | 0.17 | 0.16 | 0.16 | 0.06 | 0.05 | 0.13 | 0.11 | 0.20 |
| Allobatesnidicola | MXS | 0.94 | 0.93 | 0.93 | 0.94 | 0.97 | 1.88 | 0.03 | 0.86 | 0.12 | 0.14 | 0.14 | 0.12 | 0.07 | 0.25 | 0.11 | 0.33 |
| Allobatesnidicola | RDF | 0.78 | 0.70 | 0.70 | 0.70 | 0.82 | 1.40 | 0.30 | 0.46 | 0.09 | 0.11 | 0.11 | 0.11 | 0.06 | 0.21 | 0.11 | 0.28 |
| Allobatesnidicola | SVM | 0.52 | 0.47 | 0.47 | 0.67 | 0.79 | 1.33 | 0.13 | 0.33 | 0.40 | 0.42 | 0.42 | 0.19 | 0.13 | 0.38 | 0.17 | 0.56 |
| Allobatesnidicola | Ensemble | 0.83 | 0.67 | 0.67 | 0.76 | 0.86 | 1.52 | 0.03 | 0.55 | 0.13 | 0.22 | 0.22 | 0.14 | 0.09 | 0.28 | 0.11 | 0.32 |

| | | | | | | | | | | | | | | | | | |
|---|---|---|---|---|---|---|---|---|---|---|---|---|---|---|---|---|---|
| Allobatesniputidea | MXS | 1.00 | 0.98 | 0.98 | 0.98 | 0.99 | 1.96 | 0.00 | 0.93 | 0.01 | 0.05 | 0.05 | 0.05 | 0.02 | 0.09 | 0.00 | 0.19 |
| Allobatesniputidea | RDF | 0.97 | 0.94 | 0.94 | 0.94 | 0.97 | 1.88 | 0.06 | 0.65 | 0.03 | 0.07 | 0.07 | 0.07 | 0.04 | 0.13 | 0.07 | 0.38 |
| Allobatesniputidea | SVM | 0.99 | 0.98 | 0.98 | 0.98 | 0.99 | 1.95 | 0.03 | 0.89 | 0.03 | 0.05 | 0.05 | 0.05 | 0.03 | 0.11 | 0.05 | 0.24 |
| Allobatesniputidea | Ensemble | 0.99 | 0.93 | 0.93 | 0.93 | 0.96 | 1.86 | 0.01 | 0.70 | 0.01 | 0.06 | 0.06 | 0.06 | 0.03 | 0.12 | 0.04 | 0.32 |
| Allobatespaleovarzensis | MXS | 0.99 | 0.98 | 0.98 | 0.98 | 0.99 | 1.96 | 0.00 | 0.94 | 0.02 | 0.08 | 0.08 | 0.06 | 0.04 | 0.13 | 0.00 | 0.19 |
| Allobatespaleovarzensis | RDF | 1.00 | 1.00 | 1.00 | 1.00 | 1.00 | 2.00 | 0.00 | 1.00 | 0.00 | 0.00 | 0.00 | 0.00 | 0.00 | 0.00 | 0.00 | 0.00 |
| Allobatespaleovarzensis | SVM | 1.00 | 1.00 | 1.00 | 1.00 | 1.00 | 2.00 | 0.00 | 1.00 | 0.00 | 0.00 | 0.00 | 0.00 | 0.00 | 0.00 | 0.00 | 0.00 |
| Allobatespaleovarzensis | Ensemble | 1.00 | 1.00 | 1.00 | 1.00 | 1.00 | 2.00 | 0.00 | 1.00 | 0.00 | 0.00 | 0.00 | 0.00 | 0.00 | 0.00 | 0.00 | 0.00 |
| Allobatessumtuosus | MXS | 1.00 | 1.00 | 1.00 | 1.00 | 1.00 | 2.00 | 0.00 | 1.00 | 0.00 | 0.00 | 0.00 | 0.00 | 0.00 | 0.00 | 0.00 | 0.00 |
| Allobatessumtuosus | RDF | 0.68 | 0.35 | 0.35 | 0.65 | 0.77 | 1.30 | 0.00 | 0.80 | 0.21 | 0.41 | 0.41 | 0.20 | 0.13 | 0.40 | 0.00 | 0.27 |
| Allobatessumtuosus | SVM | 0.60 | 0.60 | 0.60 | 0.80 | 0.87 | 1.60 | 0.00 | 1.00 | 0.52 | 0.52 | 0.52 | 0.26 | 0.17 | 0.52 | 0.00 | 0.00 |
| Allobatessumtuosus | Ensemble | 0.80 | 0.65 | 0.65 | 0.82 | 0.88 | 1.63 | 0.00 | 0.81 | 0.28 | 0.47 | 0.47 | 0.24 | 0.16 | 0.48 | 0.00 | 0.37 |
| Allobatestapajos | MXS | 0.74 | 0.67 | 0.67 | 0.75 | 0.86 | 1.50 | 0.00 | 0.80 | 0.05 | 0.00 | 0.00 | 0.00 | 0.00 | 0.00 | 0.00 | 0.26 |
| Allobatestapajos | RDF | 0.57 | 0.40 | 0.40 | 0.61 | 0.76 | 1.23 | 0.07 | 0.83 | 0.14 | 0.14 | 0.14 | 0.03 | 0.02 | 0.06 | 0.14 | 0.28 |
| Allobatestapajos | SVM | 0.62 | 0.47 | 0.47 | 0.67 | 0.79 | 1.33 | 0.03 | 0.66 | 0.25 | 0.28 | 0.28 | 0.14 | 0.09 | 0.28 | 0.11 | 0.33 |
| Allobatestapajos | Ensemble | 0.63 | 0.37 | 0.37 | 0.62 | 0.76 | 1.23 | 0.00 | 0.39 | 0.07 | 0.11 | 0.11 | 0.05 | 0.03 | 0.09 | 0.00 | 0.24 |
| Allobatestrilineatus | MXS | 0.71 | 0.47 | 0.47 | 0.63 | 0.77 | 1.26 | 0.09 | 0.17 | 0.05 | 0.07 | 0.07 | 0.02 | 0.02 | 0.05 | 0.10 | 0.20 |
| Allobatestrilineatus | RDF | 0.87 | 0.67 | 0.67 | 0.74 | 0.85 | 1.48 | 0.08 | 0.37 | 0.06 | 0.11 | 0.11 | 0.07 | 0.04 | 0.13 | 0.08 | 0.18 |
| Allobatestrilineatus | SVM | 0.86 | 0.66 | 0.66 | 0.72 | 0.83 | 1.44 | 0.14 | 0.48 | 0.06 | 0.08 | 0.08 | 0.05 | 0.04 | 0.11 | 0.10 | 0.21 |
| Allobatestrilineatus | Ensemble | 0.85 | 0.68 | 0.68 | 0.74 | 0.85 | 1.48 | 0.10 | 0.44 | 0.04 | 0.08 | 0.08 | 0.05 | 0.03 | 0.10 | 0.07 | 0.18 |
| Allobateszaparo | MXS | 0.93 | 0.84 | 0.84 | 0.86 | 0.92 | 1.72 | 0.08 | 0.43 | 0.02 | 0.06 | 0.06 | 0.04 | 0.03 | 0.09 | 0.05 | 0.20 |
| Allobateszaparo | RDF | 0.94 | 0.91 | 0.91 | 0.91 | 0.95 | 1.83 | 0.08 | 0.55 | 0.06 | 0.09 | 0.09 | 0.08 | 0.05 | 0.17 | 0.07 | 0.42 |
| Allobateszaparo | SVM | 0.96 | 0.92 | 0.92 | 0.92 | 0.96 | 1.85 | 0.07 | 0.63 | 0.04 | 0.07 | 0.07 | 0.07 | 0.04 | 0.14 | 0.06 | 0.33 |
| Allobateszaparo | Ensemble | 0.94 | 0.92 | 0.92 | 0.92 | 0.96 | 1.84 | 0.08 | 0.59 | 0.05 | 0.07 | 0.07 | 0.07 | 0.04 | 0.15 | 0.07 | 0.36 |
| Ameeregabassleri | MXS | 0.98 | 0.93 | 0.93 | 0.94 | 0.97 | 1.88 | 0.00 | 0.79 | 0.03 | 0.12 | 0.12 | 0.10 | 0.05 | 0.19 | 0.00 | 0.33 |
| Ameeregabassleri | RDF | 0.83 | 0.65 | 0.65 | 0.77 | 0.86 | 1.53 | 0.08 | 0.48 | 0.16 | 0.36 | 0.36 | 0.17 | 0.11 | 0.34 | 0.12 | 0.37 |
| Ameeregabassleri | SVM | 0.87 | 0.75 | 0.75 | 0.81 | 0.88 | 1.62 | 0.13 | 0.66 | 0.14 | 0.31 | 0.31 | 0.18 | 0.12 | 0.36 | 0.13 | 0.30 |
| Ameeregabassleri | Ensemble | 0.91 | 0.80 | 0.80 | 0.84 | 0.91 | 1.68 | 0.08 | 0.68 | 0.10 | 0.20 | 0.20 | 0.14 | 0.09 | 0.29 | 0.12 | 0.36 |
| Ameeregabilinguis | MXS | 0.99 | 0.93 | 0.93 | 0.94 | 0.97 | 1.88 | 0.00 | 0.61 | 0.01 | 0.05 | 0.05 | 0.05 | 0.02 | 0.09 | 0.00 | 0.31 |
| Ameeregabilinguis | RDF | 0.92 | 0.77 | 0.77 | 0.80 | 0.89 | 1.60 | 0.07 | 0.38 | 0.04 | 0.09 | 0.09 | 0.07 | 0.05 | 0.15 | 0.09 | 0.14 |
| Ameeregabilinguis | SVM | 0.97 | 0.87 | 0.87 | 0.88 | 0.93 | 1.76 | 0.05 | 0.51 | 0.03 | 0.08 | 0.08 | 0.07 | 0.04 | 0.14 | 0.07 | 0.26 |
| Ameeregabilinguis | Ensemble | 0.97 | 0.88 | 0.88 | 0.89 | 0.94 | 1.77 | 0.06 | 0.56 | 0.03 | 0.09 | 0.09 | 0.08 | 0.05 | 0.16 | 0.07 | 0.26 |
| Ameeregabraccata | MXS | 0.78 | 0.67 | 0.67 | 0.72 | 0.83 | 1.44 | 0.17 | 0.55 | 0.12 | 0.16 | 0.16 | 0.11 | 0.07 | 0.22 | 0.18 | 0.33 |

| | | | | | | | | | | | | | | | | | |
|---|---|---|---|---|---|---|---|---|---|---|---|---|---|---|---|---|---|
| Ameeregabraccata | RDF | 0.92 | 0.77 | 0.77 | 0.83 | 0.90 | 1.65 | 0.00 | 0.69 | 0.05 | 0.16 | 0.16 | 0.12 | 0.07 | 0.24 | 0.00 | 0.27 |
| Ameeregabraccata | SVM | 0.81 | 0.70 | 0.70 | 0.76 | 0.86 | 1.52 | 0.07 | 0.65 | 0.09 | 0.11 | 0.11 | 0.09 | 0.05 | 0.18 | 0.14 | 0.31 |
| Ameeregabraccata | Ensemble | 0.83 | 0.67 | 0.67 | 0.75 | 0.86 | 1.50 | 0.00 | 0.79 | 0.06 | 0.00 | 0.00 | 0.00 | 0.00 | 0.00 | 0.00 | 0.28 |
| Ameeregaflavopicta | MXS | 0.93 | 0.82 | 0.82 | 0.87 | 0.93 | 1.74 | 0.00 | 0.70 | 0.10 | 0.22 | 0.22 | 0.15 | 0.09 | 0.29 | 0.00 | 0.32 |
| Ameeregaflavopicta | RDF | 0.89 | 0.76 | 0.76 | 0.82 | 0.89 | 1.64 | 0.08 | 0.58 | 0.13 | 0.26 | 0.26 | 0.17 | 0.11 | 0.35 | 0.14 | 0.36 |
| Ameeregaflavopicta | SVM | 0.84 | 0.70 | 0.70 | 0.76 | 0.85 | 1.52 | 0.12 | 0.48 | 0.16 | 0.25 | 0.25 | 0.18 | 0.12 | 0.37 | 0.19 | 0.42 |
| Ameeregaflavopicta | Ensemble | 0.89 | 0.78 | 0.78 | 0.83 | 0.90 | 1.66 | 0.08 | 0.57 | 0.14 | 0.27 | 0.27 | 0.19 | 0.12 | 0.38 | 0.14 | 0.49 |
| Ameeregahahneli | MXS | 0.82 | 0.56 | 0.56 | 0.67 | 0.80 | 1.34 | 0.10 | 0.68 | 0.03 | 0.11 | 0.11 | 0.05 | 0.04 | 0.11 | 0.09 | 0.08 |
| Ameeregahahneli | RDF | 0.96 | 0.85 | 0.85 | 0.86 | 0.93 | 1.73 | 0.04 | 0.69 | 0.01 | 0.05 | 0.05 | 0.04 | 0.02 | 0.08 | 0.03 | 0.16 |
| Ameeregahahneli | SVM | 0.96 | 0.83 | 0.83 | 0.85 | 0.92 | 1.70 | 0.07 | 0.63 | 0.02 | 0.08 | 0.08 | 0.06 | 0.03 | 0.11 | 0.05 | 0.14 |
| Ameeregahahneli | Ensemble | 0.96 | 0.85 | 0.85 | 0.86 | 0.93 | 1.73 | 0.07 | 0.77 | 0.02 | 0.05 | 0.05 | 0.05 | 0.03 | 0.09 | 0.05 | 0.09 |
| Ameeregamacero | MXS | 0.86 | 0.83 | 0.83 | 0.85 | 0.91 | 1.69 | 0.13 | 0.83 | 0.15 | 0.21 | 0.21 | 0.17 | 0.11 | 0.34 | 0.13 | 0.22 |
| Ameeregamacero | RDF | 0.88 | 0.70 | 0.70 | 0.76 | 0.86 | 1.52 | 0.05 | 0.44 | 0.06 | 0.11 | 0.11 | 0.07 | 0.05 | 0.14 | 0.11 | 0.09 |
| Ameeregamacero | SVM | 0.93 | 0.85 | 0.85 | 0.87 | 0.92 | 1.73 | 0.08 | 0.68 | 0.08 | 0.13 | 0.13 | 0.12 | 0.07 | 0.24 | 0.12 | 0.31 |
| Ameeregamacero | Ensemble | 0.93 | 0.85 | 0.85 | 0.86 | 0.92 | 1.72 | 0.10 | 0.62 | 0.07 | 0.13 | 0.13 | 0.12 | 0.07 | 0.24 | 0.13 | 0.33 |
| Ameeregaparvula | MXS | 0.91 | 0.77 | 0.77 | 0.79 | 0.88 | 1.59 | 0.11 | 0.66 | 0.02 | 0.05 | 0.05 | 0.04 | 0.03 | 0.09 | 0.04 | 0.11 |
| Ameeregaparvula | RDF | 0.99 | 0.94 | 0.94 | 0.95 | 0.97 | 1.89 | 0.00 | 0.54 | 0.00 | 0.02 | 0.02 | 0.02 | 0.01 | 0.04 | 0.00 | 0.25 |
| Ameeregaparvula | SVM | 0.98 | 0.87 | 0.87 | 0.88 | 0.93 | 1.76 | 0.06 | 0.47 | 0.01 | 0.06 | 0.06 | 0.06 | 0.03 | 0.11 | 0.06 | 0.24 |
| Ameeregaparvula | Ensemble | 0.99 | 0.90 | 0.90 | 0.91 | 0.95 | 1.82 | 0.02 | 0.31 | 0.01 | 0.04 | 0.04 | 0.04 | 0.02 | 0.07 | 0.03 | 0.19 |
| Ameeregapetersi | MXS | 0.70 | 0.50 | 0.50 | 0.66 | 0.79 | 1.32 | 0.08 | 0.30 | 0.12 | 0.20 | 0.20 | 0.08 | 0.06 | 0.17 | 0.12 | 0.29 |
| Ameeregapetersi | RDF | 0.78 | 0.60 | 0.60 | 0.71 | 0.82 | 1.41 | 0.10 | 0.36 | 0.14 | 0.21 | 0.21 | 0.13 | 0.09 | 0.27 | 0.13 | 0.28 |
| Ameeregapetersi | SVM | 0.71 | 0.60 | 0.60 | 0.73 | 0.84 | 1.46 | 0.00 | 0.49 | 0.25 | 0.24 | 0.24 | 0.10 | 0.07 | 0.20 | 0.00 | 0.58 |
| Ameeregapetersi | Ensemble | 0.81 | 0.60 | 0.60 | 0.75 | 0.84 | 1.49 | 0.03 | 0.45 | 0.16 | 0.34 | 0.34 | 0.17 | 0.11 | 0.34 | 0.08 | 0.33 |
| Ameeregapicta | MXS | 0.94 | 0.84 | 0.84 | 0.86 | 0.92 | 1.72 | 0.02 | 0.25 | 0.04 | 0.06 | 0.06 | 0.04 | 0.03 | 0.09 | 0.04 | 0.20 |
| Ameeregapicta | RDF | 0.95 | 0.85 | 0.85 | 0.87 | 0.93 | 1.73 | 0.08 | 0.52 | 0.04 | 0.10 | 0.10 | 0.08 | 0.05 | 0.17 | 0.08 | 0.32 |
| Ameeregapicta | SVM | 0.97 | 0.87 | 0.87 | 0.89 | 0.94 | 1.78 | 0.01 | 0.40 | 0.02 | 0.05 | 0.05 | 0.04 | 0.02 | 0.07 | 0.03 | 0.11 |
| Ameeregapicta | Ensemble | 0.97 | 0.88 | 0.88 | 0.90 | 0.95 | 1.80 | 0.01 | 0.57 | 0.03 | 0.10 | 0.10 | 0.07 | 0.04 | 0.15 | 0.03 | 0.24 |
| nomaloglossusbaeobatrach | MXS | 0.97 | 0.81 | 0.81 | 0.84 | 0.91 | 1.68 | 0.02 | 0.59 | 0.01 | 0.03 | 0.03 | 0.02 | 0.01 | 0.04 | 0.03 | 0.27 |
| nomaloglossusbaeobatrach | RDF | 0.99 | 0.92 | 0.92 | 0.93 | 0.96 | 1.86 | 0.02 | 0.58 | 0.00 | 0.03 | 0.03 | 0.03 | 0.02 | 0.06 | 0.02 | 0.18 |
| nomaloglossusbaeobatrach | SVM | 1.00 | 0.97 | 0.97 | 0.97 | 0.98 | 1.94 | 0.02 | 0.52 | 0.00 | 0.02 | 0.02 | 0.02 | 0.01 | 0.03 | 0.02 | 0.27 |
| nomaloglossusbaeobatrach | Ensemble | 1.00 | 0.96 | 0.96 | 0.96 | 0.98 | 1.93 | 0.02 | 0.72 | 0.00 | 0.03 | 0.03 | 0.02 | 0.01 | 0.05 | 0.02 | 0.25 |
| Anomaloglossusbeebei | MXS | 1.00 | 1.00 | 1.00 | 1.00 | 1.00 | 2.00 | 0.00 | 1.00 | 0.00 | 0.00 | 0.00 | 0.00 | 0.00 | 0.00 | 0.00 | 0.00 |
| Anomaloglossusbeebei | RDF | 0.82 | 0.58 | 0.58 | 0.70 | 0.82 | 1.41 | 0.06 | 0.31 | 0.09 | 0.18 | 0.18 | 0.08 | 0.05 | 0.15 | 0.10 | 0.17 |

| | | | | | | | | | | | | | | | | | |
|---|---|---|---|---|---|---|---|---|---|---|---|---|---|---|---|---|---|
| Anomaloglossusbeebei | SVM | 0.92 | 0.82 | 0.82 | 0.85 | 0.91 | 1.70 | 0.08 | 0.55 | 0.09 | 0.18 | 0.18 | 0.12 | 0.07 | 0.24 | 0.10 | 0.34 |
| Anomaloglossusbeebei | Ensemble | 0.89 | 0.82 | 0.82 | 0.84 | 0.91 | 1.69 | 0.12 | 0.49 | 0.11 | 0.18 | 0.18 | 0.12 | 0.07 | 0.24 | 0.10 | 0.36 |
| Anomaloglossusdegranville | MXS | 0.87 | 0.64 | 0.64 | 0.70 | 0.82 | 1.41 | 0.15 | 0.54 | 0.04 | 0.11 | 0.11 | 0.08 | 0.05 | 0.15 | 0.09 | 0.19 |
| Anomaloglossusdegranville | RDF | 0.89 | 0.74 | 0.74 | 0.78 | 0.88 | 1.56 | 0.10 | 0.58 | 0.04 | 0.08 | 0.08 | 0.05 | 0.03 | 0.10 | 0.06 | 0.09 |
| Anomaloglossusdegranville | SVM | 0.95 | 0.80 | 0.80 | 0.83 | 0.90 | 1.65 | 0.06 | 0.45 | 0.04 | 0.09 | 0.09 | 0.07 | 0.04 | 0.14 | 0.07 | 0.20 |
| Anomaloglossusdegranville | Ensemble | 0.93 | 0.78 | 0.78 | 0.80 | 0.89 | 1.61 | 0.09 | 0.61 | 0.03 | 0.06 | 0.06 | 0.05 | 0.03 | 0.09 | 0.05 | 0.18 |
| Anomaloglossuskaiei | MXS | 0.88 | 0.85 | 0.85 | 0.92 | 0.95 | 1.83 | 0.00 | 1.00 | 0.32 | 0.34 | 0.34 | 0.18 | 0.12 | 0.36 | 0.00 | 0.00 |
| Anomaloglossuskaiei | RDF | 0.78 | 0.65 | 0.65 | 0.77 | 0.86 | 1.53 | 0.00 | 0.83 | 0.18 | 0.24 | 0.24 | 0.16 | 0.10 | 0.32 | 0.00 | 0.25 |
| Anomaloglossuskaiei | SVM | 0.98 | 0.95 | 0.95 | 0.97 | 0.98 | 1.93 | 0.00 | 0.95 | 0.08 | 0.16 | 0.16 | 0.11 | 0.06 | 0.21 | 0.00 | 0.16 |
| Anomaloglossuskaiei | Ensemble | 0.98 | 0.95 | 0.95 | 0.97 | 0.98 | 1.93 | 0.00 | 0.94 | 0.08 | 0.16 | 0.16 | 0.11 | 0.06 | 0.21 | 0.00 | 0.17 |
| Anomaloglossuspraderioi | MXS | 0.83 | 0.65 | 0.65 | 0.78 | 0.87 | 1.57 | 0.00 | 0.83 | 0.17 | 0.34 | 0.34 | 0.19 | 0.12 | 0.39 | 0.00 | 0.25 |
| Anomaloglossuspraderioi | RDF | 0.58 | 0.30 | 0.30 | 0.65 | 0.77 | 1.30 | 0.00 | 0.75 | 0.33 | 0.48 | 0.48 | 0.24 | 0.16 | 0.48 | 0.00 | 0.50 |
| Anomaloglossuspraderioi | SVM | 0.70 | 0.55 | 0.55 | 0.70 | 0.82 | 1.40 | 0.00 | 0.95 | 0.16 | 0.16 | 0.16 | 0.11 | 0.06 | 0.21 | 0.00 | 0.16 |
| Anomaloglossuspraderioi | Ensemble | 0.75 | 0.60 | 0.60 | 0.75 | 0.85 | 1.50 | 0.00 | 1.00 | 0.29 | 0.32 | 0.32 | 0.18 | 0.11 | 0.36 | 0.00 | 0.00 |
| Anomaloglossusstepheni | MXS | 0.96 | 0.88 | 0.88 | 0.90 | 0.94 | 1.79 | 0.03 | 0.71 | 0.04 | 0.13 | 0.13 | 0.11 | 0.06 | 0.22 | 0.08 | 0.31 |
| Anomaloglossusstepheni | RDF | 0.90 | 0.80 | 0.80 | 0.83 | 0.90 | 1.66 | 0.08 | 0.59 | 0.10 | 0.16 | 0.16 | 0.12 | 0.07 | 0.25 | 0.12 | 0.32 |
| Anomaloglossusstepheni | SVM | 0.77 | 0.73 | 0.73 | 0.78 | 0.87 | 1.55 | 0.18 | 0.36 | 0.28 | 0.28 | 0.28 | 0.14 | 0.09 | 0.28 | 0.12 | 0.41 |
| Anomaloglossusstepheni | Ensemble | 0.95 | 0.88 | 0.88 | 0.89 | 0.94 | 1.78 | 0.05 | 0.69 | 0.06 | 0.13 | 0.13 | 0.12 | 0.07 | 0.23 | 0.11 | 0.34 |
| Dendrobatesleucomelas | MXS | 0.87 | 0.69 | 0.69 | 0.76 | 0.86 | 1.51 | 0.03 | 0.52 | 0.02 | 0.04 | 0.04 | 0.02 | 0.01 | 0.04 | 0.03 | 0.15 |
| Dendrobatesleucomelas | RDF | 0.99 | 0.95 | 0.95 | 0.96 | 0.98 | 1.91 | 0.00 | 0.57 | 0.00 | 0.01 | 0.01 | 0.01 | 0.00 | 0.02 | 0.01 | 0.17 |
| Dendrobatesleucomelas | SVM | 0.97 | 0.85 | 0.85 | 0.86 | 0.93 | 1.73 | 0.07 | 0.75 | 0.01 | 0.04 | 0.04 | 0.03 | 0.02 | 0.06 | 0.04 | 0.08 |
| Dendrobatesleucomelas | Ensemble | 0.97 | 0.91 | 0.91 | 0.92 | 0.96 | 1.83 | 0.04 | 0.79 | 0.01 | 0.03 | 0.03 | 0.03 | 0.02 | 0.06 | 0.03 | 0.14 |
| Dendrobatestinctorius | MXS | 0.95 | 0.77 | 0.77 | 0.79 | 0.88 | 1.58 | 0.14 | 0.86 | 0.02 | 0.06 | 0.06 | 0.05 | 0.03 | 0.10 | 0.08 | 0.14 |
| Dendrobatestinctorius | RDF | 1.00 | 0.97 | 0.97 | 0.97 | 0.98 | 1.93 | 0.02 | 0.74 | 0.01 | 0.04 | 0.04 | 0.04 | 0.02 | 0.08 | 0.03 | 0.23 |
| Dendrobatestinctorius | SVM | 0.99 | 0.97 | 0.97 | 0.97 | 0.98 | 1.94 | 0.02 | 0.63 | 0.00 | 0.02 | 0.02 | 0.02 | 0.01 | 0.04 | 0.02 | 0.18 |
| Dendrobatestinctorius | Ensemble | 1.00 | 0.98 | 0.98 | 0.98 | 0.99 | 1.95 | 0.01 | 0.83 | 0.00 | 0.03 | 0.03 | 0.02 | 0.01 | 0.05 | 0.02 | 0.18 |
| Hyloxalusnexipus | MXS | 0.98 | 0.90 | 0.90 | 0.91 | 0.95 | 1.83 | 0.00 | 0.53 | 0.02 | 0.07 | 0.07 | 0.06 | 0.03 | 0.12 | 0.00 | 0.33 |
| Hyloxalusnexipus | RDF | 1.00 | 0.97 | 0.97 | 0.98 | 0.99 | 1.95 | 0.00 | 0.87 | 0.01 | 0.06 | 0.06 | 0.05 | 0.03 | 0.11 | 0.00 | 0.27 |
| Hyloxalusnexipus | SVM | 1.00 | 0.99 | 0.99 | 0.99 | 0.99 | 1.98 | 0.00 | 0.93 | 0.01 | 0.05 | 0.05 | 0.04 | 0.02 | 0.08 | 0.00 | 0.22 |
| Hyloxalusnexipus | Ensemble | 1.00 | 0.97 | 0.97 | 0.98 | 0.99 | 1.95 | 0.00 | 0.86 | 0.01 | 0.06 | 0.06 | 0.05 | 0.03 | 0.11 | 0.00 | 0.29 |
| Hyloxalussauli | MXS | 0.97 | 0.92 | 0.92 | 0.93 | 0.96 | 1.85 | 0.04 | 0.73 | 0.05 | 0.10 | 0.10 | 0.10 | 0.05 | 0.19 | 0.08 | 0.36 |
| Hyloxalussauli | RDF | 0.95 | 0.86 | 0.86 | 0.89 | 0.94 | 1.78 | 0.00 | 0.67 | 0.06 | 0.13 | 0.13 | 0.10 | 0.06 | 0.21 | 0.00 | 0.35 |
| Hyloxalussauli | SVM | 0.98 | 0.92 | 0.92 | 0.93 | 0.96 | 1.87 | 0.00 | 0.76 | 0.04 | 0.10 | 0.10 | 0.09 | 0.05 | 0.17 | 0.00 | 0.31 |

| | | | | | | | | | | | | | | | | | |
|---|---|---|---|---|---|---|---|---|---|---|---|---|---|---|---|---|---|
| Hyloxalussauli | Ensemble | 0.99 | 0.94 | 0.94 | 0.95 | 0.97 | 1.90 | 0.00 | 0.84 | 0.02 | 0.10 | 0.10 | 0.08 | 0.04 | 0.16 | 0.00 | 0.26 |
| Ranitomeyaamazonica | MXS | 0.97 | 0.85 | 0.85 | 0.86 | 0.92 | 1.72 | 0.06 | 0.87 | 0.02 | 0.05 | 0.05 | 0.04 | 0.03 | 0.09 | 0.06 | 0.16 |
| Ranitomeyaamazonica | RDF | 0.99 | 0.92 | 0.92 | 0.92 | 0.96 | 1.84 | 0.05 | 0.55 | 0.01 | 0.04 | 0.04 | 0.04 | 0.02 | 0.07 | 0.03 | 0.23 |
| Ranitomeyaamazonica | SVM | 0.99 | 0.93 | 0.93 | 0.93 | 0.96 | 1.86 | 0.04 | 0.78 | 0.01 | 0.04 | 0.04 | 0.04 | 0.02 | 0.07 | 0.03 | 0.13 |
| Ranitomeyaamazonica | Ensemble | 0.99 | 0.93 | 0.93 | 0.93 | 0.97 | 1.87 | 0.04 | 0.80 | 0.01 | 0.04 | 0.04 | 0.04 | 0.02 | 0.08 | 0.03 | 0.16 |
| Ranitomeyafantastica | MXS | 0.82 | 0.62 | 0.62 | 0.73 | 0.84 | 1.45 | 0.00 | 0.36 | 0.04 | 0.06 | 0.06 | 0.04 | 0.02 | 0.08 | 0.00 | 0.16 |
| Ranitomeyafantastica | RDF | 0.92 | 0.76 | 0.76 | 0.80 | 0.89 | 1.61 | 0.04 | 0.44 | 0.06 | 0.13 | 0.13 | 0.10 | 0.06 | 0.20 | 0.08 | 0.23 |
| Ranitomeyafantastica | SVM | 0.92 | 0.90 | 0.90 | 0.90 | 0.95 | 1.81 | 0.08 | 0.82 | 0.10 | 0.11 | 0.11 | 0.10 | 0.06 | 0.20 | 0.10 | 0.24 |
| Ranitomeyafantastica | Ensemble | 0.90 | 0.74 | 0.74 | 0.79 | 0.88 | 1.57 | 0.10 | 0.43 | 0.07 | 0.16 | 0.16 | 0.10 | 0.06 | 0.21 | 0.11 | 0.23 |
| Ranitomeyaimitator | MXS | 0.98 | 0.88 | 0.88 | 0.89 | 0.94 | 1.79 | 0.03 | 0.44 | 0.01 | 0.04 | 0.04 | 0.03 | 0.02 | 0.06 | 0.04 | 0.20 |
| Ranitomeyaimitator | RDF | 0.94 | 0.78 | 0.78 | 0.81 | 0.90 | 1.63 | 0.03 | 0.38 | 0.03 | 0.08 | 0.08 | 0.06 | 0.04 | 0.12 | 0.06 | 0.27 |
| Ranitomeyaimitator | SVM | 0.96 | 0.91 | 0.91 | 0.92 | 0.96 | 1.83 | 0.00 | 0.46 | 0.02 | 0.03 | 0.03 | 0.02 | 0.01 | 0.04 | 0.00 | 0.09 |
| Ranitomeyaimitator | Ensemble | 0.98 | 0.88 | 0.88 | 0.90 | 0.94 | 1.79 | 0.03 | 0.44 | 0.02 | 0.06 | 0.06 | 0.04 | 0.02 | 0.09 | 0.04 | 0.21 |
| Ranitomeyareticulata | MXS | 0.97 | 0.93 | 0.93 | 0.93 | 0.96 | 1.86 | 0.06 | 0.71 | 0.04 | 0.07 | 0.07 | 0.07 | 0.04 | 0.13 | 0.07 | 0.27 |
| Ranitomeyareticulata | RDF | 0.96 | 0.93 | 0.93 | 0.93 | 0.96 | 1.86 | 0.07 | 0.55 | 0.05 | 0.07 | 0.07 | 0.07 | 0.04 | 0.13 | 0.07 | 0.41 |
| Ranitomeyareticulata | SVM | 0.95 | 0.93 | 0.93 | 0.93 | 0.96 | 1.86 | 0.07 | 0.65 | 0.05 | 0.07 | 0.07 | 0.07 | 0.04 | 0.13 | 0.07 | 0.33 |
| Ranitomeyareticulata | Ensemble | 0.96 | 0.93 | 0.93 | 0.93 | 0.96 | 1.86 | 0.07 | 0.68 | 0.05 | 0.07 | 0.07 | 0.07 | 0.04 | 0.13 | 0.07 | 0.28 |
| Ranitomeyasirensis | MXS | 0.82 | 0.59 | 0.59 | 0.66 | 0.80 | 1.33 | 0.21 | 0.36 | 0.04 | 0.11 | 0.11 | 0.06 | 0.04 | 0.12 | 0.08 | 0.21 |
| Ranitomeyasirensis | RDF | 0.95 | 0.89 | 0.89 | 0.90 | 0.94 | 1.79 | 0.09 | 0.56 | 0.03 | 0.06 | 0.06 | 0.06 | 0.03 | 0.12 | 0.05 | 0.22 |
| Ranitomeyasirensis | SVM | 0.92 | 0.76 | 0.76 | 0.79 | 0.88 | 1.58 | 0.12 | 0.51 | 0.02 | 0.06 | 0.06 | 0.04 | 0.03 | 0.09 | 0.07 | 0.14 |
| Ranitomeyasirensis | Ensemble | 0.93 | 0.82 | 0.82 | 0.83 | 0.90 | 1.66 | 0.12 | 0.59 | 0.03 | 0.09 | 0.09 | 0.08 | 0.05 | 0.16 | 0.07 | 0.18 |
| Ranitomeyatoraro | MXS | 0.88 | 0.78 | 0.78 | 0.82 | 0.90 | 1.63 | 0.03 | 0.49 | 0.06 | 0.09 | 0.09 | 0.07 | 0.04 | 0.13 | 0.05 | 0.19 |
| Ranitomeyatoraro | RDF | 0.93 | 0.87 | 0.87 | 0.87 | 0.93 | 1.74 | 0.12 | 0.58 | 0.07 | 0.11 | 0.11 | 0.11 | 0.07 | 0.23 | 0.11 | 0.37 |
| Ranitomeyatoraro | SVM | 0.97 | 0.90 | 0.90 | 0.91 | 0.95 | 1.82 | 0.02 | 0.58 | 0.03 | 0.06 | 0.06 | 0.06 | 0.03 | 0.11 | 0.05 | 0.31 |
| Ranitomeyatoraro | Ensemble | 0.94 | 0.84 | 0.84 | 0.86 | 0.92 | 1.72 | 0.06 | 0.56 | 0.05 | 0.09 | 0.09 | 0.08 | 0.04 | 0.15 | 0.08 | 0.22 |
| Ranitomeyauakarii | MXS | 0.87 | 0.72 | 0.72 | 0.74 | 0.85 | 1.48 | 0.19 | 0.55 | 0.04 | 0.06 | 0.06 | 0.05 | 0.04 | 0.11 | 0.08 | 0.23 |
| Ranitomeyauakarii | RDF | 0.94 | 0.76 | 0.76 | 0.79 | 0.88 | 1.59 | 0.11 | 0.40 | 0.04 | 0.11 | 0.11 | 0.08 | 0.05 | 0.16 | 0.08 | 0.22 |
| Ranitomeyauakarii | SVM | 0.95 | 0.78 | 0.78 | 0.81 | 0.89 | 1.62 | 0.10 | 0.53 | 0.03 | 0.11 | 0.11 | 0.09 | 0.05 | 0.18 | 0.10 | 0.16 |
| Ranitomeyauakarii | Ensemble | 0.94 | 0.78 | 0.78 | 0.81 | 0.89 | 1.62 | 0.12 | 0.50 | 0.03 | 0.10 | 0.10 | 0.08 | 0.05 | 0.16 | 0.07 | 0.05 |
| Ranitomeyavanzolinii | MXS | 0.74 | 0.60 | 0.60 | 0.72 | 0.84 | 1.44 | 0.00 | 0.69 | 0.17 | 0.14 | 0.14 | 0.06 | 0.05 | 0.13 | 0.00 | 0.29 |
| Ranitomeyavanzolinii | RDF | 0.78 | 0.73 | 0.73 | 0.80 | 0.89 | 1.60 | 0.00 | 0.92 | 0.14 | 0.14 | 0.14 | 0.11 | 0.06 | 0.21 | 0.00 | 0.25 |
| Ranitomeyavanzolinii | SVM | 0.34 | 0.20 | 0.20 | 0.57 | 0.72 | 1.14 | 0.00 | 0.36 | 0.21 | 0.28 | 0.28 | 0.10 | 0.08 | 0.21 | 0.00 | 0.46 |
| Ranitomeyavanzolinii | Ensemble | 0.77 | 0.67 | 0.67 | 0.71 | 0.83 | 1.42 | 0.17 | 0.59 | 0.08 | 0.00 | 0.00 | 0.04 | 0.03 | 0.09 | 0.18 | 0.30 |

| | | | | | | | | | | | | | | | | |
|---|---|---|---|---|---|---|---|---|---|---|---|---|---|---|---|---|
| Ranitomeyavariabilis | MXS | 0.93 | 0.81 | 0.81 | 0.84 | 0.91 | 1.68 | 0.02 | 0.61 | 0.01 | 0.02 | 0.02 | 0.02 | 0.01 | 0.03 | 0.01 | 0.20 |
| Ranitomeyavariabilis | RDF | 0.99 | 0.94 | 0.94 | 0.94 | 0.97 | 1.89 | 0.02 | 0.69 | 0.01 | 0.03 | 0.03 | 0.03 | 0.02 | 0.06 | 0.02 | 0.19 |
| Ranitomeyavariabilis | SVM | 0.98 | 0.92 | 0.92 | 0.92 | 0.96 | 1.84 | 0.02 | 0.67 | 0.01 | 0.03 | 0.03 | 0.03 | 0.02 | 0.06 | 0.03 | 0.09 |
| Ranitomeyavariabilis | Ensemble | 0.99 | 0.92 | 0.92 | 0.92 | 0.96 | 1.84 | 0.04 | 0.67 | 0.01 | 0.03 | 0.03 | 0.03 | 0.02 | 0.06 | 0.03 | 0.13 |
| Ranitomeyaventrimaculata | MXS | 0.87 | 0.74 | 0.74 | 0.80 | 0.88 | 1.60 | 0.04 | 0.50 | 0.08 | 0.18 | 0.18 | 0.12 | 0.07 | 0.24 | 0.07 | 0.30 |
| Ranitomeyaventrimaculata | RDF | 0.93 | 0.81 | 0.81 | 0.83 | 0.91 | 1.67 | 0.09 | 0.44 | 0.05 | 0.10 | 0.10 | 0.08 | 0.05 | 0.16 | 0.10 | 0.21 |
| Ranitomeyaventrimaculata | SVM | 0.93 | 0.83 | 0.83 | 0.85 | 0.92 | 1.70 | 0.04 | 0.49 | 0.04 | 0.09 | 0.09 | 0.07 | 0.04 | 0.14 | 0.07 | 0.27 |
| Ranitomeyaventrimaculata | Ensemble | 0.93 | 0.81 | 0.81 | 0.84 | 0.91 | 1.67 | 0.10 | 0.49 | 0.06 | 0.14 | 0.14 | 0.11 | 0.06 | 0.22 | 0.10 | 0.37 |
| Allobatestalamancae | MXS | 0.92 | 0.70 | 0.70 | 0.75 | 0.85 | 1.49 | 0.15 | 0.64 | 0.03 | 0.10 | 0.10 | 0.06 | 0.04 | 0.13 | 0.07 | 0.19 |
| Allobatestalamancae | RDF | 0.95 | 0.83 | 0.83 | 0.84 | 0.91 | 1.69 | 0.09 | 0.71 | 0.03 | 0.06 | 0.06 | 0.06 | 0.03 | 0.12 | 0.07 | 0.15 |
| Allobatestalamancae | SVM | 0.97 | 0.87 | 0.87 | 0.88 | 0.94 | 1.76 | 0.05 | 0.53 | 0.02 | 0.06 | 0.06 | 0.06 | 0.03 | 0.11 | 0.05 | 0.28 |
| Allobatestalamancae | Ensemble | 0.97 | 0.87 | 0.87 | 0.88 | 0.93 | 1.75 | 0.07 | 0.73 | 0.02 | 0.05 | 0.05 | 0.04 | 0.02 | 0.08 | 0.05 | 0.09 |
| Ameerega_trivitatta | MXS | 0.95 | 0.85 | 0.85 | 0.86 | 0.92 | 1.72 | 0.12 | 0.90 | 0.02 | 0.03 | 0.03 | 0.03 | 0.02 | 0.07 | 0.05 | 0.06 |
| Ameerega_trivitatta | RDF | 0.99 | 0.92 | 0.92 | 0.92 | 0.96 | 1.84 | 0.06 | 0.63 | 0.00 | 0.03 | 0.03 | 0.02 | 0.01 | 0.05 | 0.02 | 0.12 |
| Ameerega_trivitatta | SVM | 0.99 | 0.91 | 0.91 | 0.92 | 0.96 | 1.83 | 0.05 | 0.45 | 0.01 | 0.03 | 0.03 | 0.02 | 0.01 | 0.05 | 0.04 | 0.19 |
| Ameerega_trivitatta | Ensemble | 0.99 | 0.93 | 0.93 | 0.93 | 0.96 | 1.86 | 0.05 | 0.60 | 0.01 | 0.03 | 0.03 | 0.02 | 0.01 | 0.05 | 0.02 | 0.17 |
| Andinobatesfulguritus | MXS | 0.90 | 0.80 | 0.80 | 0.83 | 0.90 | 1.66 | 0.10 | 0.67 | 0.09 | 0.19 | 0.19 | 0.15 | 0.09 | 0.30 | 0.12 | 0.30 |
| Andinobatesfulguritus | RDF | 0.80 | 0.50 | 0.50 | 0.66 | 0.79 | 1.31 | 0.05 | 0.22 | 0.08 | 0.11 | 0.11 | 0.05 | 0.04 | 0.11 | 0.11 | 0.17 |
| Andinobatesfulguritus | SVM | 0.91 | 0.87 | 0.87 | 0.87 | 0.93 | 1.75 | 0.10 | 0.66 | 0.10 | 0.13 | 0.13 | 0.12 | 0.07 | 0.25 | 0.12 | 0.35 |
| Andinobatesfulguritus | Ensemble | 0.89 | 0.82 | 0.82 | 0.83 | 0.90 | 1.66 | 0.13 | 0.66 | 0.10 | 0.15 | 0.15 | 0.14 | 0.08 | 0.27 | 0.13 | 0.27 |
| Andinobatesminutus | MXS | 0.97 | 0.91 | 0.91 | 0.91 | 0.95 | 1.82 | 0.08 | 0.63 | 0.02 | 0.06 | 0.06 | 0.06 | 0.03 | 0.12 | 0.07 | 0.24 |
| Andinobatesminutus | RDF | 0.98 | 0.94 | 0.94 | 0.94 | 0.97 | 1.88 | 0.05 | 0.75 | 0.02 | 0.07 | 0.07 | 0.07 | 0.04 | 0.14 | 0.05 | 0.30 |
| Andinobatesminutus | SVM | 0.98 | 0.89 | 0.89 | 0.90 | 0.95 | 1.80 | 0.02 | 0.45 | 0.01 | 0.06 | 0.06 | 0.05 | 0.03 | 0.09 | 0.04 | 0.22 |
| Andinobatesminutus | Ensemble | 0.99 | 0.95 | 0.95 | 0.95 | 0.97 | 1.90 | 0.03 | 0.70 | 0.01 | 0.05 | 0.05 | 0.05 | 0.02 | 0.09 | 0.04 | 0.28 |
| Colostethusfraterdanieli | MXS | 0.97 | 0.90 | 0.90 | 0.91 | 0.95 | 1.81 | 0.03 | 0.72 | 0.01 | 0.03 | 0.03 | 0.03 | 0.02 | 0.06 | 0.03 | 0.09 |
| Colostethusfraterdanieli | RDF | 0.99 | 0.95 | 0.95 | 0.95 | 0.97 | 1.89 | 0.02 | 0.60 | 0.00 | 0.02 | 0.02 | 0.02 | 0.01 | 0.04 | 0.02 | 0.14 |
| Colostethusfraterdanieli | SVM | 0.99 | 0.94 | 0.94 | 0.94 | 0.97 | 1.87 | 0.05 | 0.54 | 0.01 | 0.02 | 0.02 | 0.02 | 0.01 | 0.04 | 0.03 | 0.16 |
| Colostethusfraterdanieli | Ensemble | 0.99 | 0.94 | 0.94 | 0.94 | 0.97 | 1.89 | 0.03 | 0.59 | 0.01 | 0.02 | 0.02 | 0.02 | 0.01 | 0.03 | 0.02 | 0.21 |
| Colostethusinguinalis | MXS | 0.91 | 0.77 | 0.77 | 0.79 | 0.88 | 1.58 | 0.14 | 0.43 | 0.07 | 0.12 | 0.12 | 0.10 | 0.06 | 0.19 | 0.13 | 0.28 |
| Colostethusinguinalis | RDF | 0.94 | 0.83 | 0.83 | 0.85 | 0.92 | 1.69 | 0.08 | 0.46 | 0.04 | 0.09 | 0.09 | 0.09 | 0.05 | 0.17 | 0.11 | 0.23 |
| Colostethusinguinalis | SVM | 0.96 | 0.86 | 0.86 | 0.88 | 0.93 | 1.76 | 0.02 | 0.62 | 0.06 | 0.14 | 0.14 | 0.10 | 0.06 | 0.20 | 0.05 | 0.34 |
| Colostethusinguinalis | Ensemble | 0.96 | 0.88 | 0.88 | 0.89 | 0.94 | 1.78 | 0.06 | 0.61 | 0.05 | 0.11 | 0.11 | 0.10 | 0.06 | 0.21 | 0.11 | 0.35 |
| Colostethuspanamansis | MXS | 1.00 | 0.99 | 0.99 | 0.99 | 0.99 | 1.98 | 0.00 | 0.94 | 0.01 | 0.05 | 0.05 | 0.04 | 0.02 | 0.08 | 0.00 | 0.18 |

| | | | | | | | | | | | | | | | | | |
|---|---|---|---|---|---|---|---|---|---|---|---|---|---|---|---|---|---|
| Colostethuspanamansis | RDF | 0.98 | 0.93 | 0.93 | 0.93 | 0.96 | 1.86 | 0.07 | 0.71 | 0.02 | 0.08 | 0.08 | 0.08 | 0.04 | 0.15 | 0.08 | 0.32 |
| Colostethuspanamansis | SVM | 0.98 | 0.90 | 0.90 | 0.91 | 0.95 | 1.82 | 0.03 | 0.52 | 0.02 | 0.07 | 0.07 | 0.06 | 0.03 | 0.13 | 0.06 | 0.34 |
| Colostethuspanamansis | Ensemble | 0.99 | 0.93 | 0.93 | 0.94 | 0.97 | 1.88 | 0.00 | 0.72 | 0.01 | 0.08 | 0.08 | 0.07 | 0.04 | 0.13 | 0.00 | 0.30 |
| Dendrobatesauratus | MXS | 0.98 | 0.89 | 0.89 | 0.89 | 0.94 | 1.78 | 0.08 | 0.91 | 0.01 | 0.03 | 0.03 | 0.03 | 0.02 | 0.06 | 0.03 | 0.07 |
| Dendrobatesauratus | RDF | 0.99 | 0.96 | 0.96 | 0.96 | 0.98 | 1.93 | 0.03 | 0.64 | 0.00 | 0.02 | 0.02 | 0.02 | 0.01 | 0.03 | 0.01 | 0.16 |
| Dendrobatesauratus | SVM | 0.99 | 0.96 | 0.96 | 0.96 | 0.98 | 1.92 | 0.02 | 0.72 | 0.00 | 0.01 | 0.01 | 0.01 | 0.01 | 0.02 | 0.02 | 0.11 |
| Dendrobatesauratus | Ensemble | 0.99 | 0.96 | 0.96 | 0.96 | 0.98 | 1.93 | 0.03 | 0.66 | 0.00 | 0.01 | 0.01 | 0.01 | 0.01 | 0.02 | 0.01 | 0.18 |
| Dendrobatestruncatus | MXS | 0.95 | 0.84 | 0.84 | 0.87 | 0.93 | 1.73 | 0.00 | 0.47 | 0.02 | 0.05 | 0.05 | 0.03 | 0.02 | 0.06 | 0.00 | 0.14 |
| Dendrobatestruncatus | RDF | 0.89 | 0.67 | 0.67 | 0.76 | 0.86 | 1.53 | 0.00 | 0.36 | 0.07 | 0.17 | 0.17 | 0.11 | 0.06 | 0.21 | 0.00 | 0.28 |
| Dendrobatestruncatus | SVM | 0.94 | 0.81 | 0.81 | 0.85 | 0.92 | 1.70 | 0.00 | 0.49 | 0.05 | 0.12 | 0.12 | 0.08 | 0.05 | 0.16 | 0.00 | 0.20 |
| Dendrobatestruncatus | Ensemble | 0.94 | 0.79 | 0.79 | 0.83 | 0.91 | 1.66 | 0.00 | 0.49 | 0.04 | 0.12 | 0.12 | 0.08 | 0.05 | 0.17 | 0.00 | 0.26 |
| Epipedobatesanthonyi | MXS | 0.98 | 0.94 | 0.94 | 0.94 | 0.97 | 1.87 | 0.05 | 0.59 | 0.02 | 0.04 | 0.04 | 0.04 | 0.02 | 0.09 | 0.05 | 0.23 |
| Epipedobatesanthonyi | RDF | 0.98 | 0.90 | 0.90 | 0.91 | 0.95 | 1.81 | 0.04 | 0.54 | 0.02 | 0.04 | 0.04 | 0.03 | 0.02 | 0.07 | 0.04 | 0.16 |
| Epipedobatesanthonyi | SVM | 1.00 | 0.96 | 0.96 | 0.96 | 0.98 | 1.92 | 0.01 | 0.73 | 0.00 | 0.04 | 0.04 | 0.04 | 0.02 | 0.07 | 0.02 | 0.25 |
| Epipedobatesanthonyi | Ensemble | 0.99 | 0.96 | 0.96 | 0.96 | 0.98 | 1.92 | 0.03 | 0.70 | 0.01 | 0.04 | 0.04 | 0.04 | 0.02 | 0.08 | 0.03 | 0.28 |
| Epipedobatesdarwinwallace | MXS | 0.91 | 0.78 | 0.78 | 0.81 | 0.89 | 1.62 | 0.08 | 0.64 | 0.07 | 0.14 | 0.14 | 0.12 | 0.07 | 0.23 | 0.12 | 0.30 |
| Epipedobatesdarwinwallace | RDF | 0.79 | 0.55 | 0.55 | 0.70 | 0.82 | 1.40 | 0.08 | 0.27 | 0.13 | 0.33 | 0.33 | 0.13 | 0.09 | 0.25 | 0.12 | 0.24 |
| Epipedobatesdarwinwallace | SVM | 0.80 | 0.68 | 0.68 | 0.74 | 0.85 | 1.49 | 0.13 | 0.47 | 0.13 | 0.21 | 0.21 | 0.15 | 0.10 | 0.30 | 0.13 | 0.36 |
| Epipedobatesdarwinwallace | Ensemble | 0.85 | 0.83 | 0.83 | 0.83 | 0.90 | 1.66 | 0.15 | 0.70 | 0.11 | 0.12 | 0.12 | 0.12 | 0.07 | 0.24 | 0.13 | 0.21 |
| Epipedobatesmachalilla | MXS | 0.89 | 0.76 | 0.76 | 0.79 | 0.88 | 1.58 | 0.10 | 0.41 | 0.06 | 0.10 | 0.10 | 0.07 | 0.05 | 0.15 | 0.10 | 0.11 |
| Epipedobatesmachalilla | RDF | 0.94 | 0.86 | 0.86 | 0.87 | 0.93 | 1.74 | 0.07 | 0.57 | 0.04 | 0.10 | 0.10 | 0.08 | 0.05 | 0.16 | 0.08 | 0.24 |
| Epipedobatesmachalilla | SVM | 0.94 | 0.86 | 0.86 | 0.87 | 0.93 | 1.74 | 0.07 | 0.52 | 0.06 | 0.10 | 0.10 | 0.08 | 0.05 | 0.17 | 0.08 | 0.27 |
| Epipedobatesmachalilla | Ensemble | 0.94 | 0.83 | 0.83 | 0.85 | 0.91 | 1.69 | 0.07 | 0.42 | 0.05 | 0.09 | 0.09 | 0.08 | 0.05 | 0.16 | 0.10 | 0.26 |
| Epipedobatestricolor | MXS | 0.94 | 0.81 | 0.81 | 0.84 | 0.91 | 1.68 | 0.02 | 0.31 | 0.04 | 0.06 | 0.06 | 0.04 | 0.03 | 0.09 | 0.06 | 0.14 |
| Epipedobatestricolor | RDF | 0.98 | 0.90 | 0.90 | 0.91 | 0.95 | 1.82 | 0.01 | 0.59 | 0.01 | 0.05 | 0.05 | 0.04 | 0.02 | 0.09 | 0.03 | 0.24 |
| Epipedobatestricolor | SVM | 0.98 | 0.91 | 0.91 | 0.92 | 0.96 | 1.84 | 0.00 | 0.47 | 0.01 | 0.03 | 0.03 | 0.03 | 0.02 | 0.06 | 0.00 | 0.22 |
| Epipedobatestricolor | Ensemble | 0.98 | 0.87 | 0.87 | 0.89 | 0.94 | 1.77 | 0.00 | 0.45 | 0.01 | 0.05 | 0.05 | 0.04 | 0.02 | 0.07 | 0.00 | 0.24 |
| Hyloxalusawa | MXS | 0.97 | 0.96 | 0.96 | 0.96 | 0.98 | 1.93 | 0.04 | 0.89 | 0.05 | 0.06 | 0.06 | 0.06 | 0.03 | 0.12 | 0.06 | 0.21 |
| Hyloxalusawa | RDF | 0.96 | 0.94 | 0.94 | 0.94 | 0.97 | 1.89 | 0.04 | 0.76 | 0.05 | 0.09 | 0.09 | 0.08 | 0.04 | 0.15 | 0.06 | 0.32 |
| Hyloxalusawa | SVM | 0.95 | 0.89 | 0.89 | 0.90 | 0.94 | 1.79 | 0.08 | 0.68 | 0.06 | 0.12 | 0.12 | 0.11 | 0.06 | 0.22 | 0.09 | 0.30 |
| Hyloxalusawa | Ensemble | 0.97 | 0.95 | 0.95 | 0.95 | 0.97 | 1.90 | 0.04 | 0.83 | 0.04 | 0.06 | 0.06 | 0.06 | 0.03 | 0.13 | 0.06 | 0.27 |
| Hyloxalusinfraguttatus | MXS | 0.92 | 0.84 | 0.84 | 0.86 | 0.92 | 1.72 | 0.03 | 0.69 | 0.04 | 0.07 | 0.07 | 0.05 | 0.03 | 0.11 | 0.03 | 0.15 |
| Hyloxalusinfraguttatus | RDF | 0.93 | 0.83 | 0.83 | 0.86 | 0.92 | 1.71 | 0.03 | 0.63 | 0.02 | 0.07 | 0.07 | 0.05 | 0.03 | 0.11 | 0.03 | 0.18 |

| | | | | | | | | | | | | | | | | | |
|---|---|---|---|---|---|---|---|---|---|---|---|---|---|---|---|---|---|
| Hyloxalusinfraguttatus | SVM | 0.94 | 0.84 | 0.84 | 0.86 | 0.92 | 1.72 | 0.03 | 0.43 | 0.02 | 0.08 | 0.08 | 0.06 | 0.04 | 0.12 | 0.03 | 0.08 |
| Hyloxalusinfraguttatus | Ensemble | 0.93 | 0.84 | 0.84 | 0.86 | 0.92 | 1.72 | 0.04 | 0.59 | 0.02 | 0.07 | 0.07 | 0.06 | 0.03 | 0.12 | 0.05 | 0.13 |
| Hyloxalustoachi | MXS | 0.50 | 0.20 | 0.20 | 0.57 | 0.72 | 1.13 | 0.00 | 0.60 | 0.26 | 0.26 | 0.26 | 0.09 | 0.07 | 0.17 | 0.00 | 0.55 |
| Hyloxalustoachi | RDF | 0.30 | 0.05 | 0.05 | 0.52 | 0.68 | 1.03 | 0.00 | 0.06 | 0.20 | 0.16 | 0.16 | 0.05 | 0.04 | 0.11 | 0.00 | 0.56 |
| Hyloxalustoachi | SVM | 0.45 | 0.15 | 0.15 | 0.55 | 0.71 | 1.10 | 0.00 | 0.71 | 0.20 | 0.24 | 0.24 | 0.08 | 0.06 | 0.16 | 0.00 | 0.39 |
| Hyloxalustoachi | Ensemble | 0.35 | 0.05 | 0.05 | 0.52 | 0.68 | 1.03 | 0.00 | 0.27 | 0.21 | 0.16 | 0.16 | 0.05 | 0.04 | 0.11 | 0.00 | 0.41 |
| Oophagagranulifera | MXS | 0.94 | 0.90 | 0.90 | 0.91 | 0.95 | 1.81 | 0.03 | 0.41 | 0.04 | 0.05 | 0.05 | 0.04 | 0.02 | 0.09 | 0.05 | 0.20 |
| Oophagagranulifera | RDF | 0.97 | 0.89 | 0.89 | 0.90 | 0.95 | 1.80 | 0.06 | 0.57 | 0.02 | 0.06 | 0.06 | 0.05 | 0.03 | 0.11 | 0.05 | 0.20 |
| Oophagagranulifera | SVM | 0.95 | 0.89 | 0.89 | 0.90 | 0.95 | 1.80 | 0.03 | 0.53 | 0.04 | 0.06 | 0.06 | 0.05 | 0.03 | 0.10 | 0.05 | 0.11 |
| Oophagagranulifera | Ensemble | 0.97 | 0.90 | 0.90 | 0.91 | 0.95 | 1.81 | 0.03 | 0.54 | 0.03 | 0.05 | 0.05 | 0.04 | 0.02 | 0.09 | 0.05 | 0.20 |
| Oophagahistrionica | MXS | 0.84 | 0.65 | 0.65 | 0.74 | 0.85 | 1.47 | 0.04 | 0.51 | 0.03 | 0.04 | 0.04 | 0.02 | 0.02 | 0.05 | 0.04 | 0.26 |
| Oophagahistrionica | RDF | 0.96 | 0.84 | 0.84 | 0.86 | 0.92 | 1.71 | 0.07 | 0.66 | 0.02 | 0.07 | 0.07 | 0.06 | 0.03 | 0.11 | 0.04 | 0.18 |
| Oophagahistrionica | SVM | 0.92 | 0.73 | 0.73 | 0.78 | 0.88 | 1.56 | 0.06 | 0.66 | 0.02 | 0.07 | 0.07 | 0.04 | 0.03 | 0.09 | 0.06 | 0.14 |
| Oophagahistrionica | Ensemble | 0.93 | 0.80 | 0.80 | 0.82 | 0.90 | 1.64 | 0.07 | 0.71 | 0.02 | 0.07 | 0.07 | 0.06 | 0.03 | 0.11 | 0.04 | 0.13 |
| Oophagalehmanni | MXS | 0.92 | 0.83 | 0.83 | 0.88 | 0.93 | 1.75 | 0.00 | 0.78 | 0.09 | 0.18 | 0.18 | 0.13 | 0.08 | 0.26 | 0.00 | 0.29 |
| Oophagalehmanni | RDF | 0.84 | 0.73 | 0.73 | 0.85 | 0.90 | 1.70 | 0.00 | 0.48 | 0.27 | 0.41 | 0.41 | 0.21 | 0.14 | 0.42 | 0.00 | 0.82 |
| Oophagalehmanni | SVM | 0.99 | 0.97 | 0.97 | 0.98 | 0.99 | 1.95 | 0.00 | 0.93 | 0.04 | 0.11 | 0.11 | 0.08 | 0.05 | 0.16 | 0.00 | 0.22 |
| Oophagalehmanni | Ensemble | 0.98 | 0.93 | 0.93 | 0.95 | 0.97 | 1.90 | 0.00 | 0.94 | 0.05 | 0.14 | 0.14 | 0.11 | 0.06 | 0.21 | 0.00 | 0.20 |
| Oophagapumilio | MXS | 0.96 | 0.84 | 0.84 | 0.86 | 0.92 | 1.71 | 0.08 | 0.72 | 0.01 | 0.02 | 0.02 | 0.02 | 0.01 | 0.03 | 0.03 | 0.11 |
| Oophagapumilio | RDF | 0.99 | 0.92 | 0.92 | 0.92 | 0.96 | 1.84 | 0.06 | 0.71 | 0.01 | 0.03 | 0.03 | 0.03 | 0.01 | 0.05 | 0.03 | 0.17 |
| Oophagapumilio | SVM | 0.99 | 0.92 | 0.92 | 0.92 | 0.96 | 1.84 | 0.04 | 0.66 | 0.01 | 0.02 | 0.02 | 0.02 | 0.01 | 0.04 | 0.02 | 0.18 |
| Oophagapumilio | Ensemble | 0.99 | 0.92 | 0.92 | 0.92 | 0.96 | 1.84 | 0.05 | 0.88 | 0.01 | 0.03 | 0.03 | 0.02 | 0.01 | 0.05 | 0.02 | 0.05 |
| Oophagasylvatica | MXS | 0.99 | 0.95 | 0.95 | 0.95 | 0.97 | 1.90 | 0.04 | 0.55 | 0.01 | 0.02 | 0.02 | 0.02 | 0.01 | 0.05 | 0.04 | 0.22 |
| Oophagasylvatica | RDF | 0.99 | 0.96 | 0.96 | 0.96 | 0.98 | 1.93 | 0.03 | 0.69 | 0.01 | 0.04 | 0.04 | 0.03 | 0.02 | 0.07 | 0.03 | 0.28 |
| Oophagasylvatica | SVM | 0.99 | 0.97 | 0.97 | 0.97 | 0.99 | 1.95 | 0.02 | 0.77 | 0.01 | 0.03 | 0.03 | 0.03 | 0.01 | 0.05 | 0.03 | 0.28 |
| Oophagasylvatica | Ensemble | 0.99 | 0.97 | 0.97 | 0.97 | 0.98 | 1.94 | 0.02 | 0.71 | 0.01 | 0.03 | 0.03 | 0.03 | 0.01 | 0.05 | 0.03 | 0.28 |
| Phyllobatesaurotaenia | MXS | 0.94 | 0.87 | 0.87 | 0.88 | 0.93 | 1.76 | 0.07 | 0.64 | 0.06 | 0.13 | 0.13 | 0.11 | 0.07 | 0.23 | 0.09 | 0.31 |
| Phyllobatesaurotaenia | RDF | 0.83 | 0.62 | 0.62 | 0.71 | 0.83 | 1.43 | 0.07 | 0.54 | 0.09 | 0.14 | 0.14 | 0.09 | 0.06 | 0.17 | 0.09 | 0.20 |
| Phyllobatesaurotaenia | SVM | 0.94 | 0.85 | 0.85 | 0.87 | 0.93 | 1.73 | 0.05 | 0.53 | 0.06 | 0.09 | 0.09 | 0.08 | 0.05 | 0.17 | 0.08 | 0.29 |
| Phyllobatesaurotaenia | Ensemble | 0.91 | 0.77 | 0.77 | 0.81 | 0.89 | 1.62 | 0.07 | 0.61 | 0.07 | 0.16 | 0.16 | 0.12 | 0.07 | 0.25 | 0.09 | 0.30 |
| Phyllobateslugubris | MXS | 0.90 | 0.82 | 0.82 | 0.84 | 0.91 | 1.68 | 0.09 | 0.57 | 0.08 | 0.12 | 0.12 | 0.11 | 0.06 | 0.22 | 0.10 | 0.24 |
| Phyllobateslugubris | RDF | 0.93 | 0.77 | 0.77 | 0.80 | 0.88 | 1.59 | 0.11 | 0.42 | 0.05 | 0.13 | 0.13 | 0.10 | 0.06 | 0.21 | 0.13 | 0.25 |
| Phyllobateslugubris | SVM | 0.97 | 0.89 | 0.89 | 0.90 | 0.95 | 1.80 | 0.04 | 0.69 | 0.03 | 0.10 | 0.10 | 0.09 | 0.05 | 0.19 | 0.08 | 0.31 |

| | | | | | | | | | | | | | | | | |
|---|---|---|---|---|---|---|---|---|---|---|---|---|---|---|---|---|
| Phyllobateslugubris | Ensemble | 0.95 | 0.84 | 0.84 | 0.87 | 0.92 | 1.73 | 0.04 | 0.60 | 0.05 | 0.12 | 0.12 | 0.10 | 0.06 | 0.21 | 0.08 | 0.29 |
| Phyllobatesterribilis | MXS | 1.00 | 1.00 | 1.00 | 1.00 | 1.00 | 2.00 | 0.00 | 1.00 | 0.00 | 0.00 | 0.00 | 0.00 | 0.00 | 0.00 | 0.00 | 0.00 |
| Phyllobatesterribilis | RDF | 1.00 | 1.00 | 1.00 | 1.00 | 1.00 | 2.00 | 0.00 | 1.00 | 0.00 | 0.00 | 0.00 | 0.00 | 0.00 | 0.00 | 0.00 | 0.00 |
| Phyllobatesterribilis | SVM | 1.00 | 1.00 | 1.00 | 1.00 | 1.00 | 2.00 | 0.00 | 1.00 | 0.00 | 0.00 | 0.00 | 0.00 | 0.00 | 0.00 | 0.00 | 0.00 |
| Phyllobatesterribilis | Ensemble | 1.00 | 1.00 | 1.00 | 1.00 | 1.00 | 2.00 | 0.00 | 1.00 | 0.00 | 0.00 | 0.00 | 0.00 | 0.00 | 0.00 | 0.00 | 0.00 |
| Phyllobatesvittatus | MXS | 1.00 | 0.96 | 0.96 | 0.97 | 0.98 | 1.93 | 0.00 | 0.89 | 0.00 | 0.04 | 0.04 | 0.04 | 0.02 | 0.07 | 0.00 | 0.21 |
| Phyllobatesvittatus | RDF | 0.98 | 0.89 | 0.89 | 0.90 | 0.95 | 1.81 | 0.00 | 0.38 | 0.01 | 0.04 | 0.04 | 0.03 | 0.02 | 0.06 | 0.00 | 0.14 |
| Phyllobatesvittatus | SVM | 1.00 | 1.00 | 1.00 | 1.00 | 1.00 | 2.00 | 0.00 | 1.00 | 0.00 | 0.00 | 0.00 | 0.00 | 0.00 | 0.00 | 0.00 | 0.00 |
| Phyllobatesvittatus | Ensemble | 1.00 | 0.99 | 0.99 | 0.99 | 0.99 | 1.97 | 0.00 | 0.89 | 0.00 | 0.03 | 0.03 | 0.03 | 0.01 | 0.06 | 0.00 | 0.23 |
| Silverstoneiaflotator | MXS | 0.99 | 0.95 | 0.95 | 0.95 | 0.98 | 1.91 | 0.04 | 0.80 | 0.01 | 0.05 | 0.05 | 0.05 | 0.03 | 0.10 | 0.03 | 0.24 |
| Silverstoneiaflotator | RDF | 1.00 | 1.00 | 1.00 | 1.00 | 1.00 | 2.00 | 0.00 | 1.00 | 0.00 | 0.00 | 0.00 | 0.00 | 0.00 | 0.00 | 0.00 | 0.00 |
| Silverstoneiaflotator | SVM | 1.00 | 0.99 | 0.99 | 0.99 | 0.99 | 1.98 | 0.00 | 0.91 | 0.00 | 0.02 | 0.02 | 0.02 | 0.01 | 0.05 | 0.00 | 0.19 |
| Silverstoneiaflotator | Ensemble | 1.00 | 1.00 | 1.00 | 1.00 | 1.00 | 2.00 | 0.00 | 1.00 | 0.00 | 0.00 | 0.00 | 0.00 | 0.00 | 0.00 | 0.00 | 0.00 |
| Silverstoneianubicola | MXS | 0.92 | 0.76 | 0.76 | 0.80 | 0.89 | 1.60 | 0.08 | 0.77 | 0.05 | 0.11 | 0.11 | 0.07 | 0.05 | 0.15 | 0.06 | 0.15 |
| Silverstoneianubicola | RDF | 0.99 | 0.94 | 0.94 | 0.95 | 0.97 | 1.89 | 0.03 | 0.69 | 0.03 | 0.09 | 0.09 | 0.07 | 0.04 | 0.15 | 0.04 | 0.28 |
| Silverstoneianubicola | SVM | 0.94 | 0.76 | 0.76 | 0.80 | 0.88 | 1.59 | 0.08 | 0.37 | 0.04 | 0.11 | 0.11 | 0.07 | 0.04 | 0.13 | 0.06 | 0.19 |
| Silverstoneianubicola | Ensemble | 0.98 | 0.92 | 0.92 | 0.93 | 0.96 | 1.86 | 0.04 | 0.64 | 0.05 | 0.10 | 0.10 | 0.10 | 0.06 | 0.20 | 0.09 | 0.31 |
